# Blockade of IKK signaling induces RIPK1-independent apoptosis in human cells

**DOI:** 10.1101/2023.06.20.545781

**Authors:** Neha M. Nataraj, Beatrice Herrmann, Sunny Shin, Igor E. Brodsky

**Author notes:** Co-corresponding authors: Igor E. Brodsky and Sunny Shin.

## Abstract

Regulated cell death in response to microbial infection plays an important role in immune defense and is triggered by pathogen disruption of essential cellular pathways. Gramnegative bacterial pathogens in the *Yersinia* genus disrupt NF-κB signaling via translocated effectors injected by a type III secretion system (T3SS), thereby preventing induction of cytokine production and antimicrobial defense. In murine models of infection, *Yersinia* blockade of NF-κB signaling triggers cell-extrinsic apoptosis through Receptor Interacting Serine-Threonine Protein Kinase 1 (RIPK1) and caspase-8, which is required for bacterial clearance and host survival. Unexpectedly, we find that human macrophages undergo apoptosis independently of RIPK1 in response to *Yersinia* or chemical blockade of IKKα/β. Instead, IKK blockade led to decreased cFLIP expression, and overexpression of cFLIP contributed to protection from IKK blockade-induced apoptosis in human macrophages. Importantly, IKK blockade also induces RIPK1 kinase-independent apoptosis in human T cells and human pancreatic cells. Altogether, our data indicate that, in contrast to murine cells, blockade of IKK activity in human cells triggers a distinct apoptosis pathway that is independent of RIPK1. These findings have implications for the contribution of RIPK1 to cell death in humans and the efficacy of RIPK1 inhibition in human diseases.

## INTRODUCTION

Regulated cell death in response to microbial infection is critical for immune defense. Innate immune cells utilize pattern recognition receptors to detect pathogen-associated molecular patterns and elicit an inflammatory response (1,2). To evade this detection, many microbial pathogens employ mechanisms to suppress immune signaling (3,4). In response, the host has evolved compensatory mechanisms to induce cell death and inflammation in response to pathogen-mediated disruption of TNFR superfamily (5,6) and TLR3/4/TRIF signaling (6,7).

Stimulation of TNFR or TLR4 induces recruitment of RIPK1 to the receptor proximal signaling complex. RIPK1 serves as a Complex 1 scaffold for downstream signaling proteins, including TAK1 and IKKα/β, which phosphorylate RIPK1 to maintain its localization (8–15). The kinases in Complex I initiate NF-κB and MAPK signaling, resulting in inflammatory cytokine production (16). However, pathogen-mediated or pharmacological blockade of key Complex I proteins, particularly TAK1 and IKKα/β, destabilizes the complex. Released RIPK1 then recruits FADD and caspase-8 to form Complex II, mediating rapid RIPK1-dependent cell death independently of NF-κB’s role in cell survival (8,10,17).

The Gram-negative bacterial genus *Yersinia* causes human diseases ranging from gastroenteritis to plague (18,19). *Yersinia* utilizes a type III secretion system (T3SS) to inject virulence factors known as *Yersinia* outer proteins (Yops) into the host cell cytosol (20,21). One of these Yops, known as YopP in *Y. enterocolitica* and YopJ in *Y. pseudotuberculosis*, potently blocks IKKβ and TAK1 in murine macrophages, resulting in RIPK1- and caspase-8-dependent cell death (14,22–35). Importantly, RIPK1 activity is critical for caspase-8-dependent cell death, restriction of bacterial loads, and host survival during *Yersinia* infection in mice (12,33).

While murine models have been vital to elucidate mechanisms of cell death and inflammation during *Yersinia* infection, there exist notable differences in gene expression, and immune responses between human and murine immune systems (36–39), indicating that mice do not fully recapitulate human responses to infection. Notably, mice lacking RIPK1 experience acute perinatal lethality, succumbing to systemic inflammation and aberrant cell death (40), whereas humans with biallelic RIPK1 deficiency are viable, although they experience immunodeficiencies and autoinflammatory disease (41,42). Clinical studies have implicated human RIPK1 in a wide range of systemic disorders and pathologies (41–46), leading to substantial interest in developing RIPK1 therapeutics, particularly for treating inflammatory diseases and cancer (46–48). However, whether human and murine cells similarly undergo RIPK1-dependent cell death in response to blockade of IKK signaling remains poorly understood.

Here, we demonstrate that in human macrophages, cell death induced by *Yersinia* or chemical IKK blockade is independent of RIPK1, indicating that regulation of apoptosis in human macrophages is distinct from mouse macrophages. Instead, our data suggest that cell death is caused by IKK blockade-mediated downregulation of cFLIP. cFLIP overexpression partially protected human macrophages from IKK blockade-induced cell death. Moreover, we found that RIPK1 activity is also dispensable for apoptosis of human Jurkat T cells and pancreatic cells following IKK blockade. Altogether, our data demonstrate that IKK blockade triggers apoptosis independently of RIPK1 in multiple human cell types. Our findings suggest that investigation of compensatory cell death pathways is warranted in the setting of therapeutics targeting RIPK1 for human disease.

## MATERIALS AND METHODS

### Cell cultures

All cell types were maintained in a humidified incubator kept at 37°C with 5% CO_2_. **Murine bone marrow-derived macrophages** (BMDMs) were isolated and differentiated as previously described (49) and replated in a 96-well tissue culture (TC)-treated plate at a concentration of 0.5×10^5^ cells/well. **Primary human monocyte-derived macrophages** (hMDMs) were differentiated as previously described (37) and replated in a 96-well TC-treated plate at 0.3×10^5^ cells/well, or a 48-well tissue culture-treated plate at 1×10^5^ cells/well. **THP-1 monocytes** (TIB-202; American Type Culture Collection) were maintained, differentiated, and replated as previously described (38) in a 96-well TC- reated plate at 0.5×10^5^ cells/well, or a 48-well TC-treated plate at 2×10^5^ cells/well. **Jurkat T cells clone E6-1** (TIB-152; American Type Culture Collection) were kindly provided by Will Bailis (University of Pennsylvania, Philadelphia). Cells were maintained in THP-1 media (38) as recommended. The day before the experiment, cells were replated in media without antibiotics in a 96-well TC-treated plate at 0.5×10^5^ cells/well. **PANC-1** pancreatic epithelial carcinoma cells (CRL-1469; American Type Culture Collection) were maintained in DMEM supplemented with 10% (vol/vol) heat-inactivated fetal bovine serum (FBS), 100 IU/mL penicillin, and 100 μg/mL streptomycin. The day before the experiment, cells were detached with trypsin-EDTA (0.25%) and replated in media without antibiotics in a 96-well TC-treated plate at 0.5×10^5^ cells/well.

### Bacterial Strains and Growth Conditions

*Yersinia* strains described in **Table 1** were grown and induced as previously described (50). All cultures were washed and resuspended in pre-warmed serum-free media prior to infection. In all experiments, control cells were mock-infected with serum-free media.

**Table 1:** Strains

| Table 1: Strains |  |
| --- | --- |
| Strain name | Reference/Source |
| IP2666 (WT *Yptb*) | From James Bliska (ref. 52) |
| IP26 ($\Delta yopJ$ *Yptb*) | From James Bliska (ref. 22) |
| IP26 pACYC184-YopP | (ref. 56) |
| JB580v (WT *Ye*) | From Virginia Miller (ref. 54 ) |
| YVM1612 (JB580v $\Delta yopP$ *Ye*) | From Virginia Miller (unpublished) |

### Stimulations and Infections

Cell media was replaced with fresh media containing inhibitors (**Table 2**) and centrifuged at 300 x g for 1 min. 1-2 h following addition of inhibitors, cells were stimulated or infected (**Table 2**). For priming conditions, cells were treated with 100 ng/mL *E. coli* LPS for 4 h before stimulation/infection. Cells were infected at a multiplicity of infection (MOI) of 20:1, unless otherwise indicated, centrifuged at 300 x g for 5 min, and incubated at 37°C. At 1 h post-infection, cells were treated with 100 μg/mL of gentamicin. At the indicated harvest timepoints, cells were centrifuged at 300 x g for 5 min prior to sample collection.

**Table 2:**
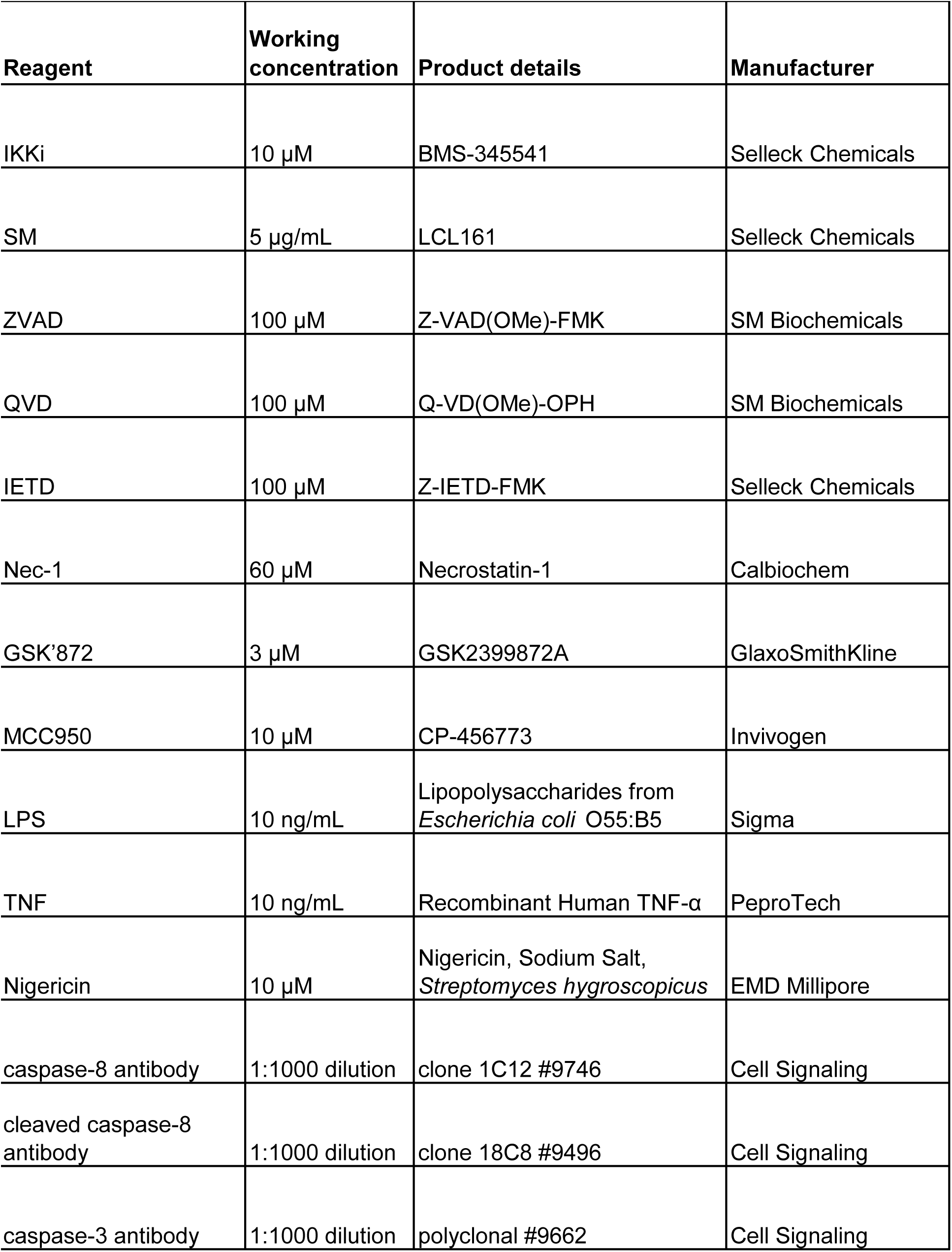
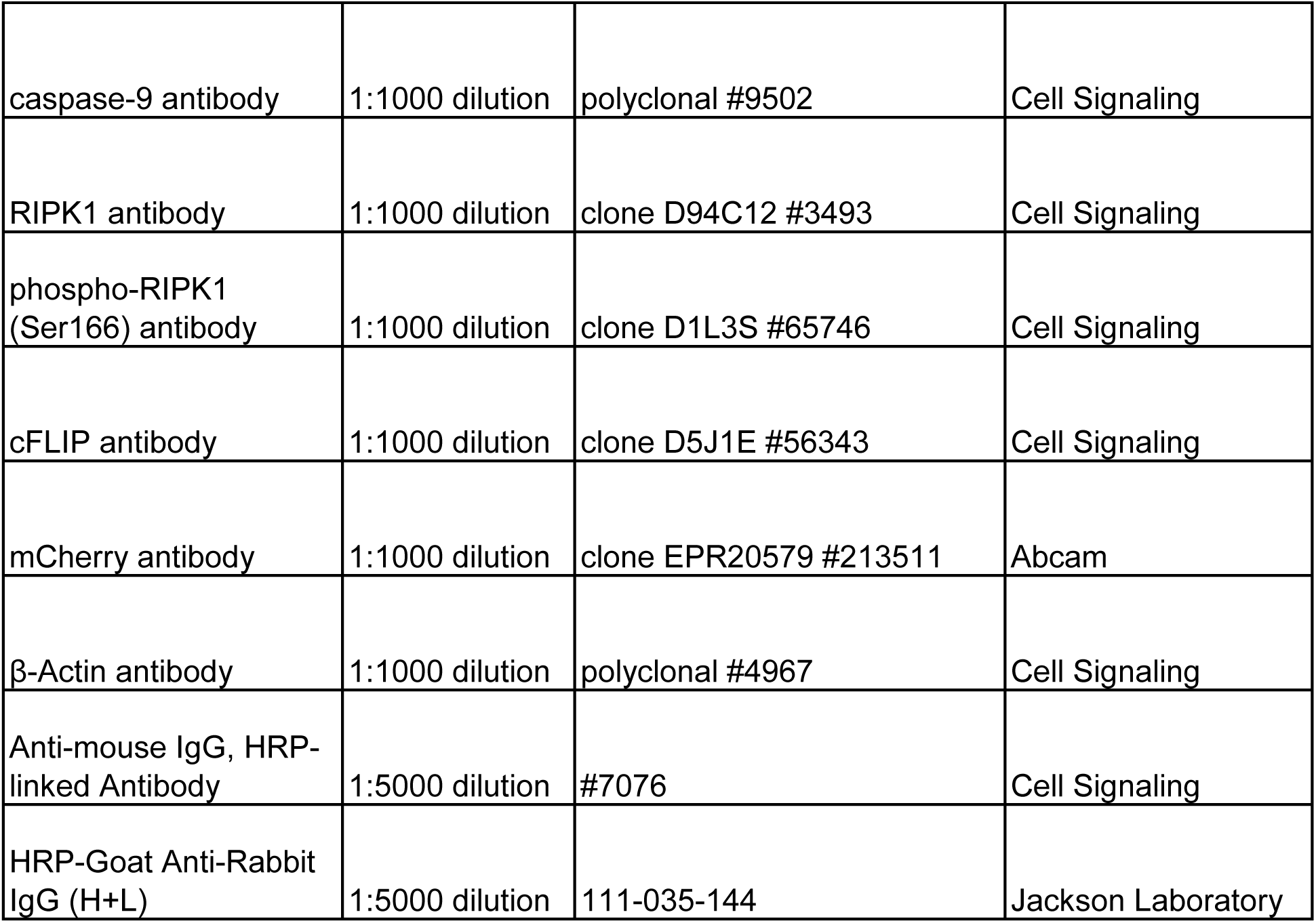
Reagents

### LDH Release

Cell supernatants were assayed for cytotoxicity by measuring loss of cellular membrane integrity via lactate dehydrogenase release, quantified using LDH Cytotoxicity Detection Kit (Sigma) according to manufacturer directions. % cytotoxicity was calculated by subtracting background (untreated/mock infected) and normalizing to maximal LDH release (1% Triton X-100).

### PI Uptake

Propidium iodide (PI) (Thermo Fisher, Waltham, MA, USA) uptake was performed as previously described (51) and detected on a BioTek Synergy HT Multi-Detection Microplate Reader (BioTek, Winooski, VT) at 485/20 excitation and 590/35 emission every 10 min for the indicated time points. % PI uptake was calculated by subtracting background (untreated) and normalizing to maximal PI uptake (1% Triton X-100).

### Cell Viability

Cells were plated in a white-walled, clear-bottom TC-treated 96 well plate and treated as indicated. ATP was detected using the CellTiter-Glo 2.0 Assay Kit (Promega) according to manufacturer instructions, and the reaction was incubated in the dark for 30-60 min. Luminescence was read on a Gen5 plate reader (BioTek). % survival was calculated by subtracting background (1% Triton X-100) and normalizing to maximal ATP levels (untreated).

### Caspase-3/7 Activity

Cells were plated in serum-free media in a white-walled, clear-bottom TC-treated 96 well plate and treated as indicated. Caspase-3/7 activity was detected using the Caspase-Glo 3/7 Assay Kit (Promega) according to manufacturer instructions. The reaction was incubated in the dark for 3 h. Luminescence was read on a Gen5 plate reader (BioTek) and values were normalized to cell signal read by ATP detection.

### Immunoblot Analysis

Cells were lysed in SDS/PAGE sample buffer (50 mM Tris-HCl pH 6.8, 2% w/v SDS, 10% v/v glycerol, 0.1% Bromophenol Blue, 2 mM DTT). Lysates were boiled and centrifuged prior to running on 4–12% Bis-Tris gels (Thermo Fisher) and transferred to PVDF membranes. Membranes were immunoblotted using primary antibodies at 1:1000 dilution and HRP-linked secondary antibodies at 1:5000 dilution (**Table 2**). Membranes were developed using Pierce ECL Plus and SuperSignal West Femto Maximum Sensitivity Substrate according to manufacturer instructions (Thermo Fisher). Uncropped blots are found in **Supplementary File**.

### Cytokine Release

Supernatants were harvested and cytokine levels were assayed using ELISA kits for human TNF and IL-18 (R&D Systems).

Generation of various THP-1 lines, qRT-PCR, and statistical analyses were performed as described in Supplementary Materials and Methods.

## RESULTS

### *Yersinia* YopP blockade of IKK signaling induces RIPK1 activity-independent apoptosis in human macrophages

*Yersinia* YopJ blockade of IKK signaling induces rapid RIPK1- and caspase-8-dependent cell death in murine bone marrow-derived macrophages (BMDMs) (12,23,31,33–35) (**Fig. S1A**). However, primary human monocyte-derived macrophages (hMDMs) infected with *Y. pseudotuberculosis* (*Yptb*) (52) did not undergo detectable cytotoxicity (**Fig. S1B**), consistent with recent reports (34). The related species *Y. entercolitica* (*Ye*) expresses a YopJ homolog, termed YopP, that is required for *Ye* to block TNF production in hMDMs (**Fig. 1A**) (22,53). In contrast to *Yptb*, *Ye* (54) induced robust YopP-dependent cell death in both hMDMs and BMDMs (**Fig. S1A, B, C**), as previously reported (55). Interestingly, a Δ*yopJ Yptb* strain expressing YopP (56) was not sufficient to recapitulate the cell death induced by WT *Ye* (**Fig. S1C**). Unexpectedly, Necrostatin-1 (Nec-1), a small molecule inhibitor of RIPK1 kinase activity, did not block *Ye-*induced death of hMDMs (**Fig. 1B, C**). In contrast, it potently blocked *Ye-* and *Yptb-*induced death of BMDMs (**Fig. S1A**), consistent with our and other published studies (31,34).

**Fig. 1.**
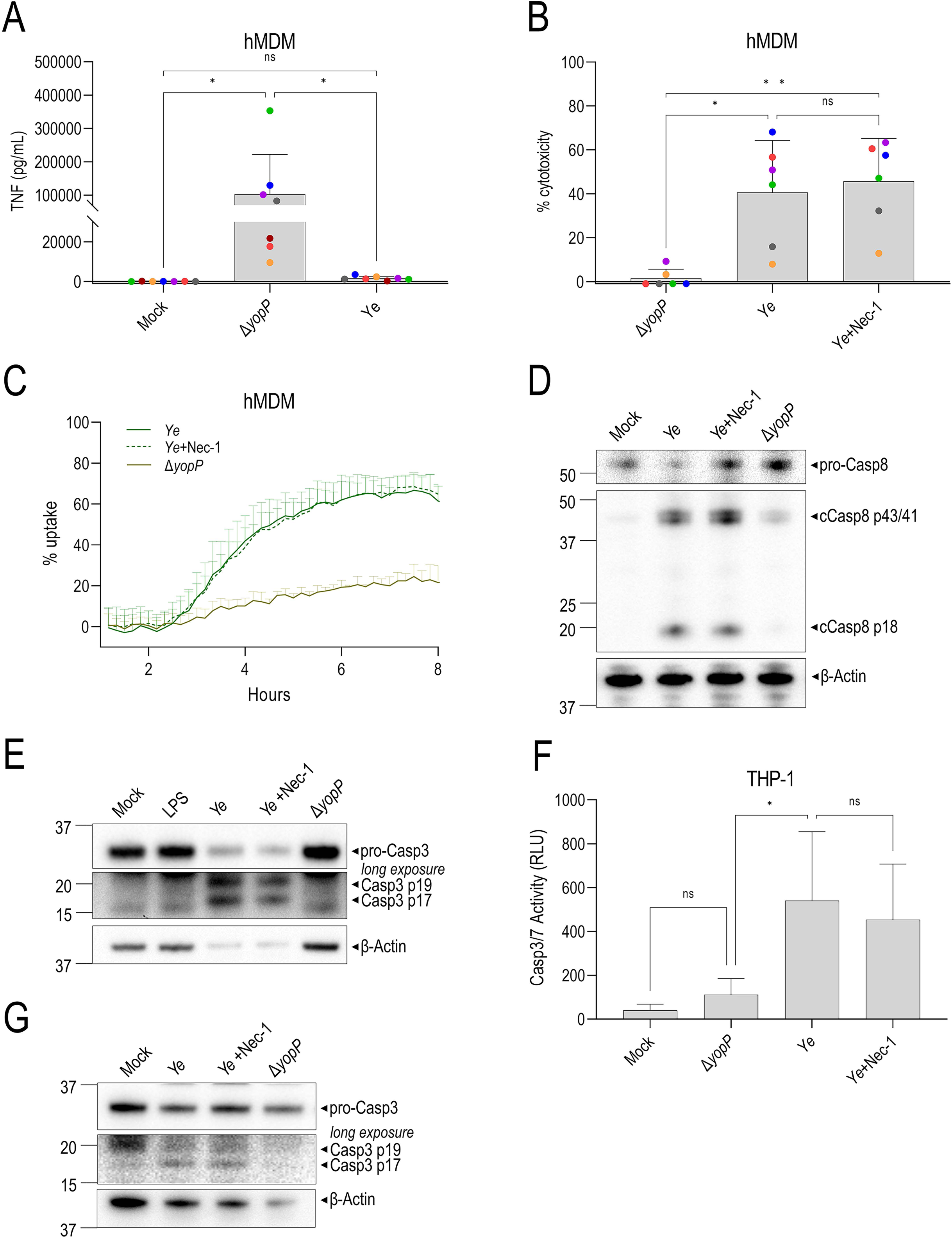
RIPK1 activity is dispensable for *Yersinia*-induced extrinsic apoptosis in human macrophages. Cells were pre-treated with Nec-1 for 1 h and mock-infected or infected with WT *Y. enterocolitica* (*Ye*) or Δ*yopP*. **(A)** TNF levels in the supernatant of hMDMs were measured by ELISA 18-19 h after infection. **(B)** Cytotoxicity was measured by lactate dehydrogenase (LDH) release from hMDMs at 18-19 h after infection and normalized to untreated cells. **(A-B**) Each data point represents the mean of triplicate wells for each of 6–7 different human donor hMDMs. **(C)** Cytotoxicity in hMDMs was measured by PI uptake in triplicate wells over 8 h. Representative of three independent experiments with different human donors. **(D-E)** Immunoblot analysis was performed on hMDM lysates 18 h after infection for caspase-8, caspase-3, and β-actin. Representative of three independent experiments. **(F)** Caspase-3/7 activity in THP-1 cells quantified 20-26 h after infection. N=5 independent experiments. **(G)** Immunoblot analysis of THP-1 lysates 24 h after infection for caspase-3 and β-actin. Representative of two independent experiments. ns, not significant, *p < 0.05, **p < 0.01, ***p < 0.001, ****p < 0.0001 by Tukey’s multiple comparisons test. Graphs depict mean + SD.

Immunoblot analysis of hMDMs demonstrated YopP-dependent caspase-8 processing into its active form that was not inhibited by Nec-1 (**Fig. 1D**), in contrast to BMDMs. *Ye* also induced YopP-dependent cleavage of the cell-intrinsic initiator caspase, caspase-9, in hMDMs (**Fig. S1D**), consistent with observations that *Yersinia* induces caspase-8-dependent cleavage of Bid, which can activate the mitochondrial apoptosis pathway (25). The downstream executioner caspase-3 was processed into the active subunits p17/p19 in a RIPK1 activity-independent manner (**Fig. 1E**). Human THP-1 macrophages also exhibited robust YopP-dependent, RIPK1 activity-independent caspase-3/7 cleavage and activation (**Fig. 1F, G**). Overall, these data demonstrate that *Ye* induces YopP-dependent, RIPK1 activity-independent apoptosis in human macrophages.

### RIPK1 activity is dispensable for IKK blockade-induced apoptosis of human macrophages

In murine BMDMs, TNF or TLR stimulation together with pharmacological TAK1 or IKK blockade induces RIPK1- and caspase-8-dependent cell death (12,34), similar to *Yersinia* infection. Consistently, hMDMs (**Fig. 2A, B**) and THP-1 cells (**Fig. 2E**) treated with LPS and an IKK inhibitor (LPS+IKKi) underwent increased cell death (**Fig. 2A, B**). However, as with *Yersinia* infection (**Fig. 1**), this cell death was not blocked by Nec-1, indicating that RIPK1 activity was not required. In murine cells, Complex I blockade induces RIPK1 autophosphorylation, which is necessary for RIPK1 to form a stable Complex II and activate caspase-8 (10,57,58). However, we did not detect RIPK1 phosphorylation during LPS+IKKi treatment in hMDMs (**Fig. 2D**).

**Fig. 2.**
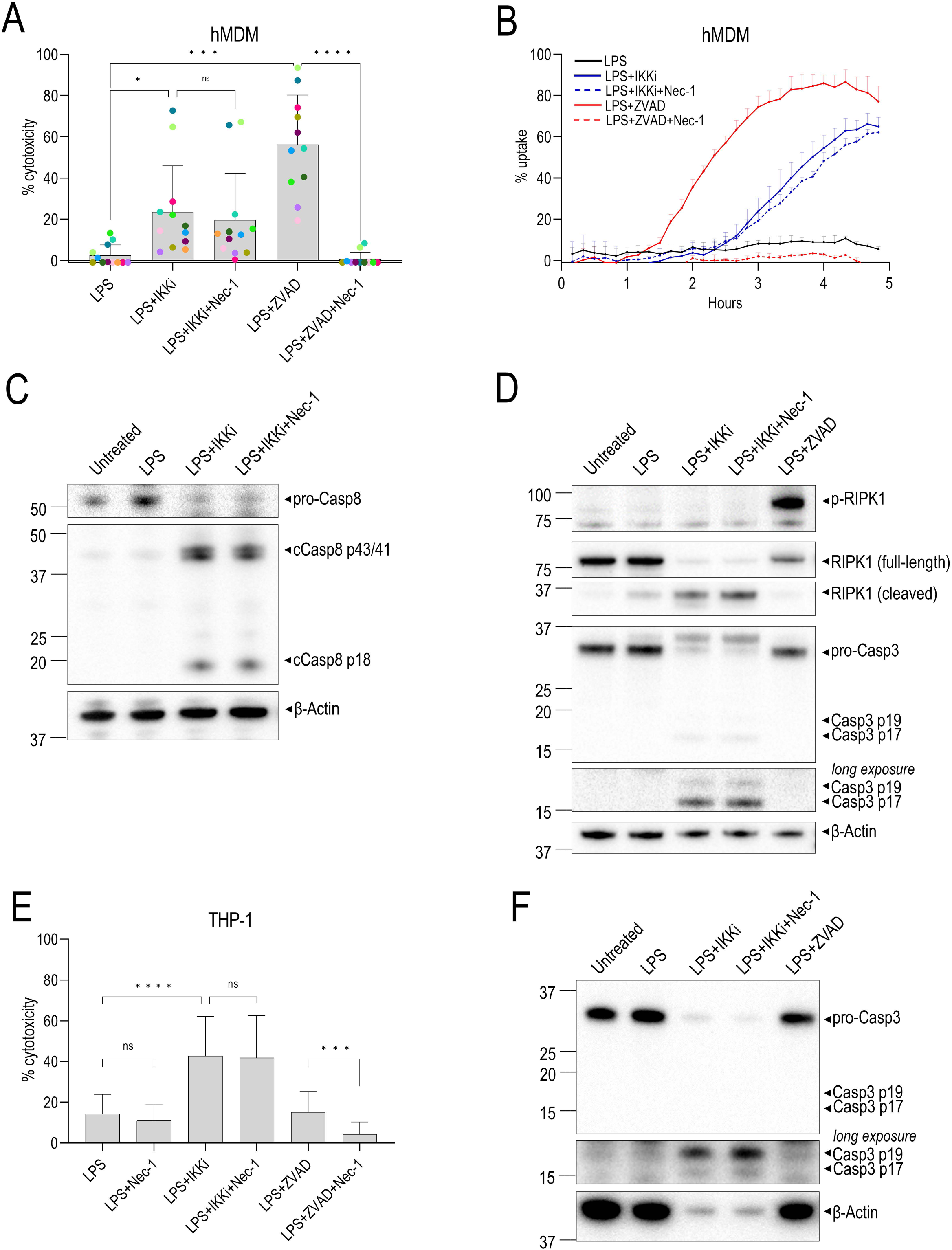
RIPK1 activity is dispensable for IKKα/β blockade-induced extrinsic apoptosis in human macrophages. Cells were pre-treated with IKKi, Nec-1, and/or ZVAD and stimulated with LPS. **(A)** Cytotoxicity in hMDMs was measured by LDH release 5-7 h post-stimulation. Each data point represents the mean of triplicate wells for 11-12 different human donors. **(B)** Cytotoxicity in hMDMs was measured by PI uptake in triplicate wells over 5 h. Representative of three independent experiments. **(C)** Immunoblot analysis was performed on hMDM lysates for caspase-8, phospho-RIPK1 (S166), RIPK1, caspase-3, and β-actin. Representative of 3-4 independent experiments. **(D)** Cytotoxicity in THP-1 cells was measured by LDH release 17-24 h post-stimulation. N=22-35. **(F)** Immunoblot analysis was performed on THP-1 lysates at 24 h for caspase-3 and β-actin. Representative of two experiments. ns, not significant, *p < 0.05, **p < 0.01, ***p < 0.001, ****p < 0.0001 by Tukey’s multiple comparisons test. Graphs depict mean + SD.

We also treated cells with LPS together with the pan-caspase inhibitor ZVAD, which engages necroptosis by blocking caspase-8 activity (59). Importantly, we observed robust RIPK1 phosphorylation (**Fig. 2D**), and Nec-1 effectively suppressed necroptosis following LPS+ZVAD treatment (**Fig. 2A, B, E**). Consistent with other studies (60), LPS+ZVAD-induced death of human macrophages was blocked by the RIPK3 inhibitor GSK’872 (**Fig. S2A, C**). In total, our data indicate that IKK blockade-induced death of human macrophages occurs independently of RIPK1 activity.

As with *Ye* infection, LPS+IKKi treatment induced processing of caspase-8 (**Fig. 2C**) and caspase-3 (**Fig. 2D, F**) into their active forms which was not blocked by Nec-1. LPS+IKKi-induced cell death was significantly reduced upon treatment with the caspase-8 inhibitor IETD and RIPK3 inhibitor GSK’872 in hMDMs (**Fig. S2B**), or with the pan-caspase inhibitor QVD and GSK’872 in THP-1 cells (**Fig. S2D**), consistent with the interpretation that LPS+IKKi induces caspase-8-dependent cell death. LPS+IKKi also induced caspase-9 cleavage (**Fig. S2E**), as observed with *Ye* infection (**Fig. S1D**).

In murine macrophages, the *Yersinia*-induced RIPK1/caspase-8 complex also mediates pyroptosis by activating the pore-forming protein Gasdermin D (GSDMD) and promoting IL-18 release (31,34,35,61). As expected, LPS+Nigericin treatment, which activates the NLRP3 inflammasome, induced IL-18 release in THP-1 cells (**Fig. S2F**). However, neither LPS+IKKi treatment nor *Yersinia* infection induced IL-18 release (**Fig. S2F**). Collectively, our data indicate that following chemical or *Yersinia-*dependent IKK blockade, human macrophages undergo caspase-8-dependent apoptosis independent of RIPK1 activity.

### RIPK1 is dispensable for IKK blockade-induced cell-extrinsic apoptosis in human macrophages

In murine cells, RIPK1 can contribute to cell-extrinsic apoptosis via both kinase-dependent and independent functions (8–12,15,57,62,63). Blockade of Complex I proteins at Checkpoint 1 occurs upstream of NF-κB-dependent gene regulation and triggers rapid RIPK1 activity-dependent cell death (7–12,17,57,64). Checkpoint 2 occurs at the level of transcriptional/translational regulation of pro-survival genes such as cFLIP and A20 (7,10,15,17,57,62). Downregulation of cFLIP leads to the loss of caspase-8-cFLIP heterodimers that normally inhibit cell death, inducing RIPK1 activity-independent cell death (6,10,17,57,63).

To genetically test the requirement for RIPK1 in IKK blockade-induced cell death, we used CRISPR/Cas9 to generate two independent *RIPK1^-/-^* THP-1 single-cell clones (**Fig. S3A-C**). *RIPK1^-/-^* THP-1 cells released significantly lower levels of TNF upon LPS stimulation (**Fig. S3D**), consistent with the reported role for RIPK1 in TLR4 signaling (65). *RIPK1^-/-^* THP-1 cells were not sensitized to LPS-induced cytotoxicity (**Fig. 3A**), similar to previous findings (42), but in contrast to prior findings with human iPSC-derived macrophages stimulated with LPS (66) and *RIPK1^-/-^* Jurkat T cells stimulated with TNF (8,17,67). Following LPS+IKKi treatment or *Ye* infection, we found that WT and *RIPK1^-/-^* THP-1 cells exhibited comparable levels of cell death (**Fig. 3A**) and caspase-3/7 activity (**Fig. 3B-C**). Consistent with the critical role of RIPK1 in necroptosis, *RIPK1^-/-^* THP-1 cells were protected from cytotoxicity in response to LPS+ZVAD (**Fig. 3A**). Altogether, these findings demonstrate that human macrophages do not require RIPK1 for LPS+IKKi-induced apoptosis.

**Fig. 3.**
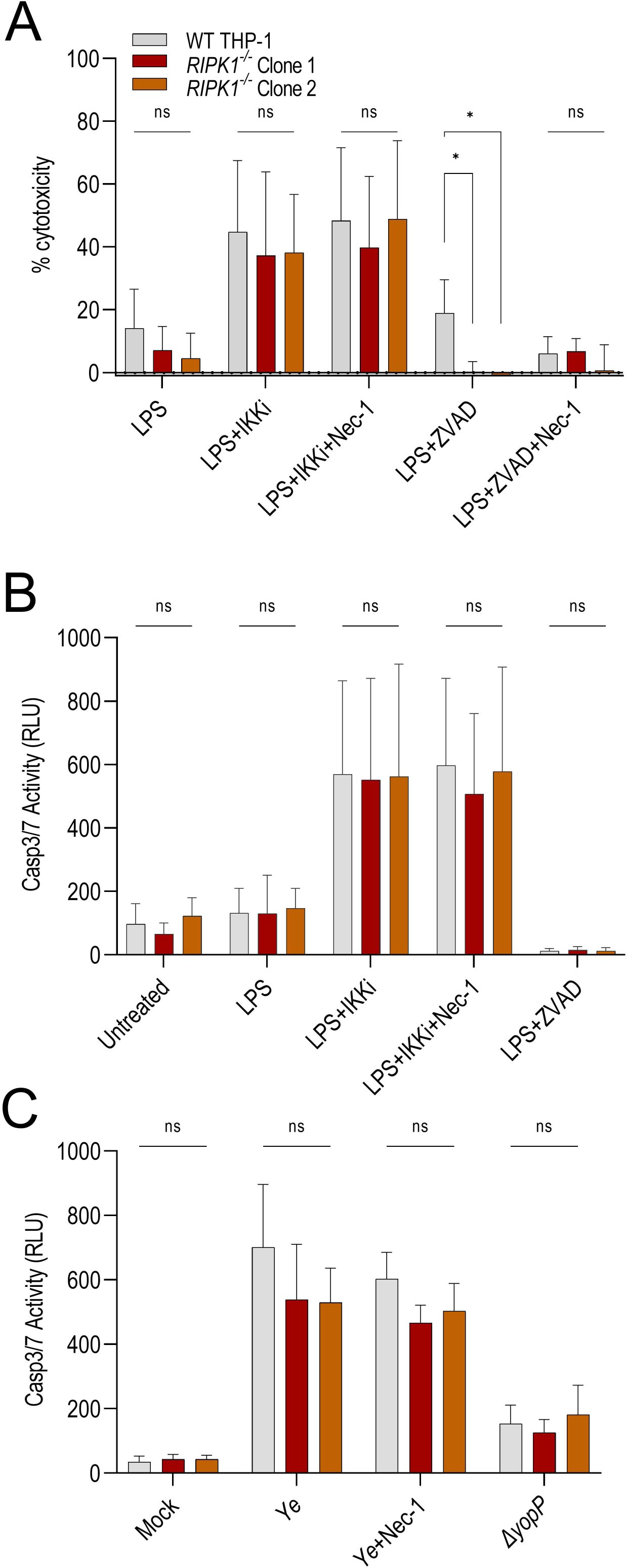
Human *RIPK1^-/-^* macrophages undergo cell-extrinsic apoptosis following IKKα/β blockade and *Yersinia* infection. WT and *RIPK1^-/-^* THP-1 macrophages were pre-treated with inhibitors and stimulated with LPS or infected with *Ye* (WT or Δ*yopP*). **(A)** Cytotoxicity was measured by LDH release 18-24 h post-stimulation. N=6-14. **(B-C)** Caspase-3/7 activity was detected and quantified by Caspase-Glo 3/7 22-24 h post-stimulation or infection. **(B)** N=3-4. **(C**) N=3. ns, not significant, *p < 0.05 by Tukey’s multiple comparisons test. Graphs depict mean + SD.

### cFLIP partially protects human macrophages from IKK blockade-induced apoptosis

Since RIPK1 was dispensable for IKK blockade-induced apoptosis in human macrophages, we hypothesized that inhibition of IKK reduces expression of pro-survival genes such as cFLIP, which limits cell-extrinsic apoptosis (17,59,61,68,69). Indeed, we observed rapid downregulation of cFLIP expression following LPS+IKKi treatment (**Fig. 4A**). We generated two independent *CFLAR^-/-^* THP-1 single cell clones (**Fig. S4A-C**) and found that they were significantly more susceptible than WT cells to caspase-3/7 activation and apoptosis following LPS or TNF stimulation alone (**Fig. 4B, C**), consistent with the established protective role of cFLIP (17,59,61,68,69). Cytotoxicity of LPS+IKKi-treated WT cells was comparable to that of *CFLAR^-/-^* cells, suggesting that IKKi-induced downregulation of cFLIP sensitizes THP-1 cells to death similarly to cells genetically lacking cFLIP (**Fig. 4B**). LPS+ZVAD treatment largely protected *CFLAR^-/-^* cells from LPS-induced apoptosis (**Fig. 4B**), consistent with our finding that *CFLAR^-/-^* cells are predisposed to undergo caspase-3/7 activation and apoptosis in response to LPS (**Fig. 4C**).

**Fig. 4.**
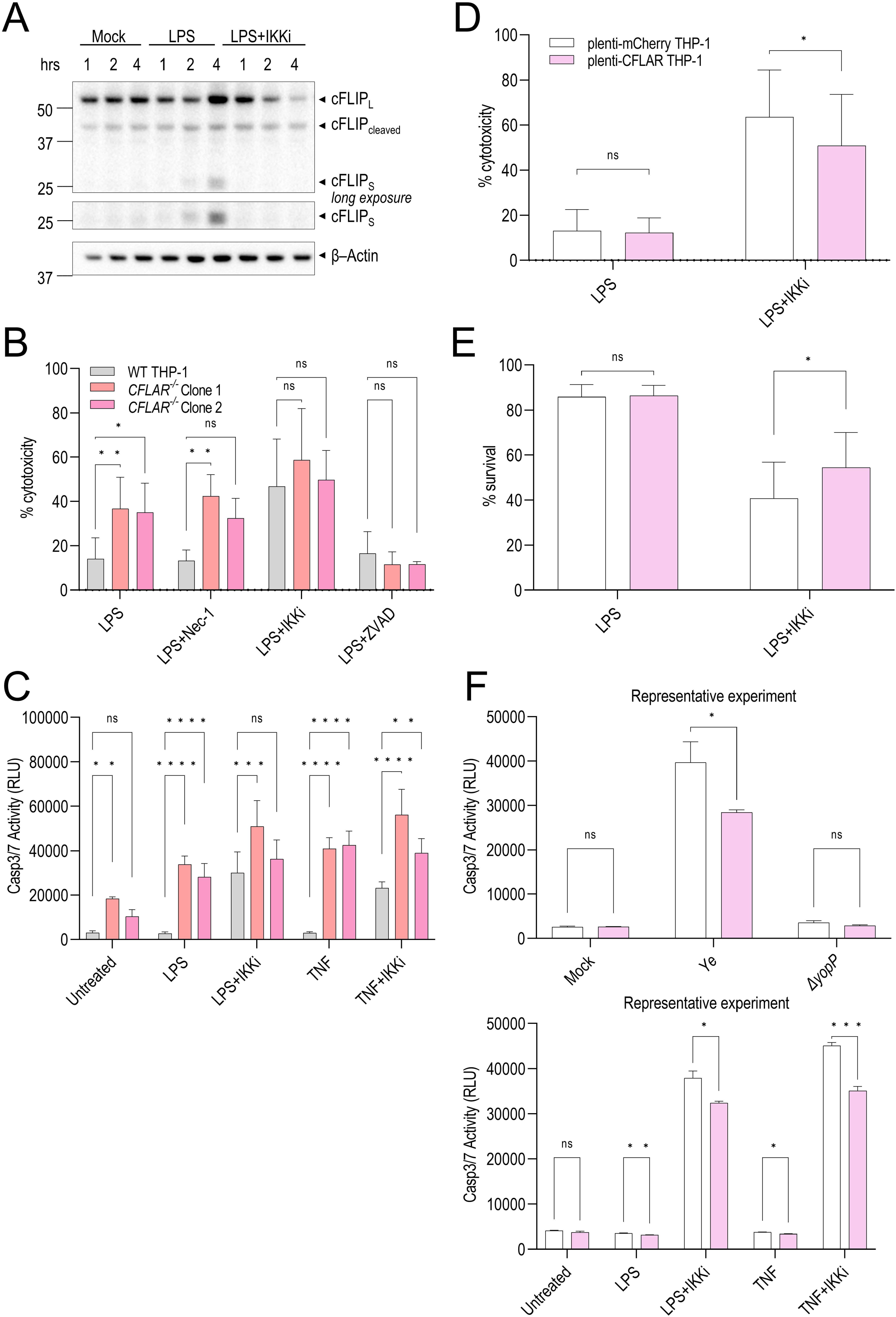
cFLIP partially regulates extrinsic apoptosis in human macrophages following IKKα/β blockade. **(A)** Immunoblot analysis was performed on WT hMDM lysates for cFLIP expression at various times post-treatment. Representative of four independent experiments**. (B-C)** WT and *CFLAR^-/-^* THP-1 macrophages were pre-treated with inhibitors and stimulated with LPS for 18-25 h. N=3-4. ns, not significant, *p < 0.05, **p < 0.01, ***p < 0.001, ****p < 0.0001 by Dunnett’s multiple comparisons test. **(B)** Cytotoxicity was measured by LDH release. **(C)** Cell viability was measured by ATP signal. **(D-F)** mCherry- and CFLAR-overexpressing THP-1 cells were pre-treated with inhibitors and stimulated with LPS/TNF or infected with *Ye* for 24 h. **(D-E)** ns, not significant, *p < 0.05 by Holm-Šídák method paired T test. **(D)** Cytotoxicity was measured by LDH release. N=5. **(E)** Cell viability was measured by ATP signal. N=4. **(F)** Caspase-3/7 activity was detected and quantified by Caspase-Glo 3/7. ns, not significant, *p < 0.05 by Holm-Šídák method unpaired T test. Representative of three independent experiments performed in triplicate. All graphs depict mean + SD.

We next tested whether cFLIP overexpression would protect human macrophages from IKK blockade-induced apoptosis by generating THP-1 cells that stably overexpressed either *CFLAR* or *mCherry* (WT control) (**Fig. S4D**). THP-1 cells overexpressing cFLIP were significantly protected from cell death following *Yersinia* infection or IKKi treatment. (**Fig. 4D-F**). However, this protection was incomplete, indicating that cFLIP is necessary but not completely sufficient to protect cells from death. Altogether, these data indicate that cFLIP overexpression partially protects against IKK blockade-induced apoptosis in human macrophages.

### RIPK1 kinase activity is dispensable for IKK blockade-induced cell death in multiple human cell types

We next sought to test whether lack of a role for RIPK1 activity in regulating cell-extrinsic apoptosis was specific to human macrophages or a feature shared by other human cells. RIPK1 was previously reported to contribute to apoptosis in both human Jurkat T cells and PANC-1 pancreatic tumor cells (17,70,71). Given the lack of TLR4 expression in Jurkat cells (72), we treated the cells instead with TNF+IKKi, which largely phenocopies LPS+IKKi-induced apoptosis (12,73). THP-1 cells and hMDMs underwent RIPK1 activity-independent cell death in response to TNF+IKKi treatment (**Fig. 5A, B**), demonstrating that IKK blockade-induced death downstream of LPS and TNF are both RIPK1 activity-independent pathways in human macrophages. Moreover, TNF+IKKi treatment in Jurkat cells (**Fig. 5C**) and PANC-1 cells (**Fig. 5D**) also induced cell death that was not blocked by Nec-1, indicating that RIPK1 activity is dispensable for cell-extrinsic apoptosis in multiple human cell types in response to IKK blockade in the setting of TLR or TNF stimulation.

**Fig. 5.**
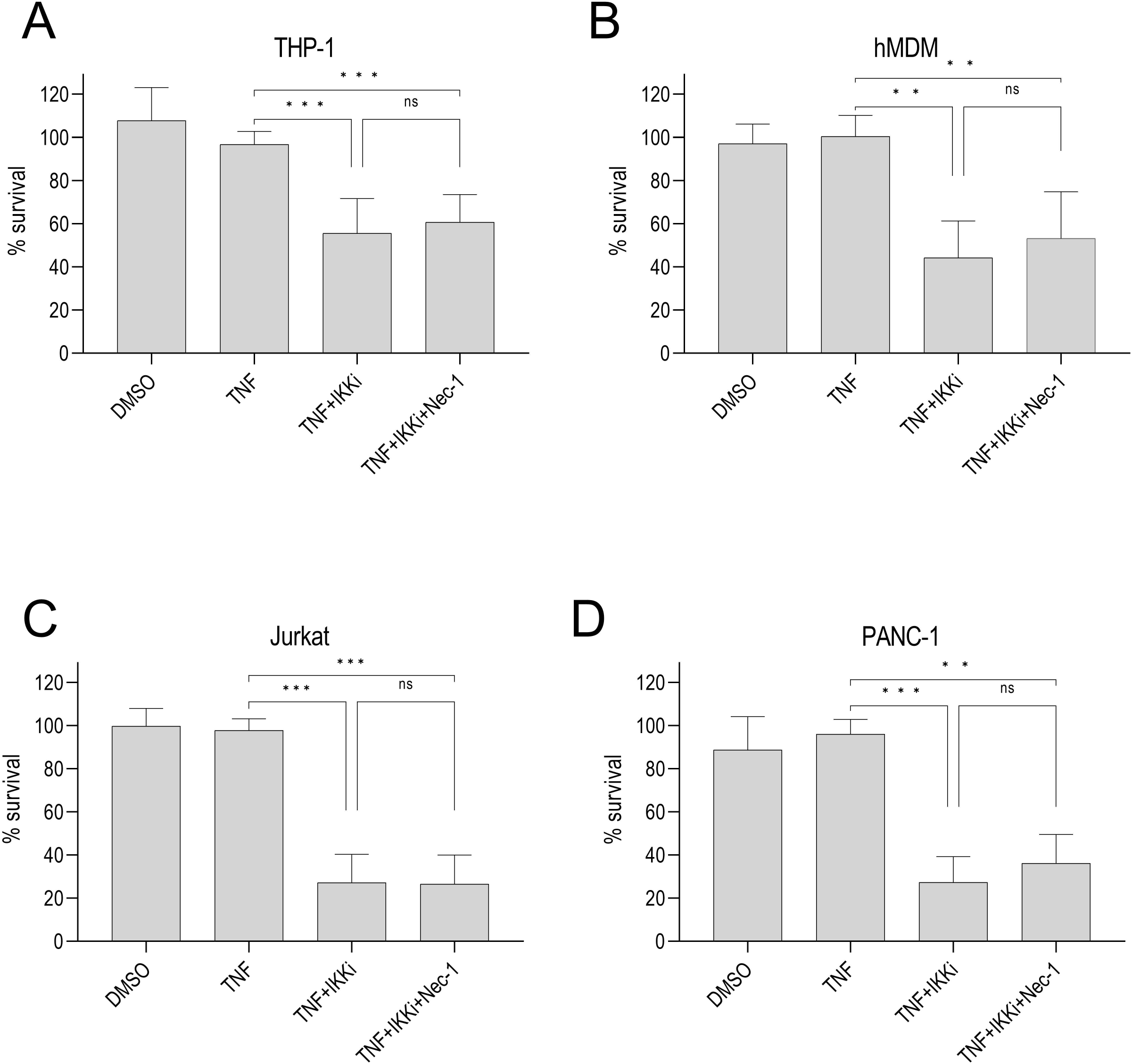
Multiple human cell types undergo RIPK1 activity-independent cell death following IKKα/β blockade. Cells were pre-treated with IKKi ± Nec-1 and stimulated with TNF. Viability was measured by ATP levels in the following cell types: **(A)** THP-1 cells, 22-25 h, N=6, **(B)** hMDMs, 5-6 h, N=4, **(C)** Jurkat cells, 12-15 h, N=3, **(D)** PANC-1 cells, 14-16 h, N=3. ns, not significant, *p < 0.05, **p < 0.01, ***p < 0.001 by Tukey’s multiple comparisons test. Graphs depict mean + SD.

The E3 ubiquitin ligases cIAP1/2 ubiquitylate RIPK1 to form a scaffold for recruitment of signaling proteins, including IKKα/β and TAK1 (8,9,57,15). SMAC mimetics (SM), which disrupt cIAPs and induce their degradation, are used extensively to induce RIPK1-dependent apoptosis. Interestingly, both THP-1 cells and hMDMs were resistant to cell death following TNF+SM treatment (**Fig. S5A, B**), consistent with previous findings that macrophages are resistant to SM-induced cell death (74). In contrast, both Jurkat T cells and PANC-1 cells were highly susceptible to cell death following TNF+SM treatment, and this was significantly reduced by Nec-1 treatment (**Fig. S5C, D**). Collectively, these data indicate that distinct factors mediate cell-extrinsic apoptosis in different human cell types in response to IKK blockade or disruption of cIAPs.

## DISCUSSION

Here, we report that unlike in murine macrophages, RIPK1 is dispensable for cell-extrinsic apoptosis in human macrophages during *Yersinia* infection or pharmacological blockade of IKK. In its role as a scaffold within Complex I, RIPK1 mediates canonical NF-κB signaling and promotes host cell survival in numerous systems (40,42,65,75). However, our data demonstrate that in the absence of RIPK1, human macrophages do not exhibit elevated susceptibility to necroptosis, in contrast to previous reports (8,17,40,41,66,67,76), and these cells can still release substantial, albeit reduced, levels of TNF (**Fig. S3D**), suggesting that RIPK1 is not entirely required for pro-inflammatory and pro-survival signaling. Furthermore, human macrophages were not susceptible to cIAP1/2 inhibition (**Fig. S5A, B**). Given that cIAP1/2 provides protection from cell death via its regulation of RIPK1, whereas cIAP1/2 inhibition was not cytotoxic to human macrophages, it is possible that pro-survival signaling complexes in human macrophages are partially RIPK1-independent. Overall, combined with our findings that IKK blockade induces RIPK1-independent cell death in human macrophages (**Fig. 3**), our studies also suggest the possibility that a RIPK1-independent checkpoint promotes pro-survival NF-κB signaling and protects human macrophages from aberrant cell death. We cannot exclude the possibility that human macrophages can undergo RIPK1-dependent apoptosis in response to other stimuli. Nonetheless, our findings demonstrate that human and murine macrophages exhibit distinct requirements for RIPK1 in IKK blockade-induced apoptosis.

PANC-1 and Jurkat cells underwent apoptosis in response to treatment with SMAC mimetic and TNF, and this death was partially RIPK1-dependent (**Fig. S5**). In contrast, upon TNF+IKKi treatment, PANC-1 and Jurkat cells underwent RIPK1 activity-independent apoptosis (**Fig. 5**). We speculate that in these cells, blocking cIAP1/2 destabilizes Complex I and releases RIPK1 to form Complex II and mediate apoptosis, whereas blocking IKKα/β does not release RIPK1 but directly targets downstream NF-κB signaling. This is in contrast to murine cells, wherein blockade of TAK1 or IKKα/β releases RIPK1 to mediate Complex II formation and apoptosis (15,34). Overall, these findings highlight the existence of cell type- and species-specific heterogeneity in regulation of cell-extrinsic apoptosis in response to blockade of immune signaling.

Interestingly, contrary to murine macrophages, we found that *Yersinia pseudotuberculosis* (*Yptb*) does not induce cell death in human macrophages. In contrast, *Y. enterocolitica* (*Ye*) induced robust cell death in human macrophages, which was dependent on YopP, a homolog of YopJ in *Yptb*. Notably, *Yptb* expressing YopP did not induce cell death (**Fig. S1C**), indicating that other Yops in *Yptb* suppress death in human macrophages. Indeed, our recent studies now suggest that *Yptb* effectors YopE/H/K synergistically suppress pyroptosis in both human IECs and macrophages, and that YopJ does not induce RIPK1-dependent apoptosis in human cells (77).

Studies examining extrinsic apoptosis following Checkpoint 1 blockade in human cells have largely investigated SMAC mimetic treatment in epithelial cell lines and report that RIPK1 mediates apoptosis in this context (13,17,42,78), which we observed as well (**Fig. S5**). Previous studies also established that RIPK1 is required for apoptosis following membrane-bound FasL treatment in human T cells (42,71,79). In contrast, our findings demonstrate that RIPK1 is not required for cell-extrinsic apoptosis following IKK blockade in human macrophages, T cells, and pancreatic epithelial cells, indicating that distinct regulatory mechanisms exist in human cells to mediate cell-extrinsic apoptosis. RIPK1 is implicated in a broad range of human diseases (43,44,46), and there is significant interest in development of therapeutics to target RIPK1 kinase activity in a variety of human inflammatory diseases, neurodegenerative conditions, and cancer metastasis (46–48). While extensive analyses in murine models highlight a key role for RIPK1 activity in promoting cell-extrinsic apoptosis and downstream responses, our results indicate that there are key differences in the contribution and function of RIPK1 activity in human disease. Our findings highlight the need to further dissect the contributions of RIPK1 to cell death in different contexts in human cells.

## ACKNOWLEDGEMENTS

We thank Drs. Virginia Miller and Kimberly Walker for generously providing WT *Y. enterocolitica* JB580v (54) and YVM1612 (JB580v *ΔyopP*) for our studies. Graphic figures were created with BioRender. This work was supported by NIH/NIAID awards R01AI139102A1 (I.E.B.), R21AI125924 (I.E.B.), R01AI118861 (S.S.), R01AI123243 (S.S.), BWF Investigators in the Pathogenesis of Infectious Disease Awards (S.S. and I.E.B.); Penn Physical Sciences Oncology Center Pilot Award (supported by NIH/NCI U54 grant CA93417) and Penn Center for Genome Integrity Pilot Award from the University of Pennsylvania; NIH/NIAID award 5T32AI141393 (N.M.N.) and AHA Predoctoral Fellowship (N.M.N).

## ETHICS STATEMENT

All studies involving primary human macrophages were performed in compliance with the requirements of the US Department of Health and Human Services and the principles expressed in the Declaration of Helsinki. Samples were obtained from the University of Pennsylvania Human Immunology Core. These samples are considered to be a secondary use of de-identified human specimens and are exempt via Title 55 Part 46, Subpart A of 46.101 (b) of the Code of Federal Regulations.

All experiments performed with mouse bone marrow-derived macrophages were approved by the Institutional Animal Care and Use Committee of the University of Pennsylvania (protocol 804523).

