## Supplemental Material for "Blockade of IKK signaling induces RIPK1-independent apoptosis in human cells"

#### SUPPLEMENTARY METHODS

##### Generation of CRISPR/Cas9 THP-1 knockout cell lines

*RIPK1*$^{-/-}$ and *CFLAR*$^{-/-}$ cells were generated using the CRISPR/Cas9 system as previously described(38). Briefly, pLentiCRISPR v2 plasmids encoding the desired guide RNA (gRNA) and Cas9 were purchased from GenScript. The following target sequences were used (5' to 3'): *RIPK1* gRNA 1: CGGCTTTCAGCACGTGCATC, *CFLAR* gRNA 3: TGCCAATGCAATCGATTATC. Production of lentiviral particles, transduction of target cells, single-cell clone selection, and sequence validation were performed as previously described(38). DNA was isolated from clones using DNeasy Blood & Tissue Kit (Qiagen, Hilden, Germany). The genomic region containing the target sequence was amplified by PCR using the following primers (all 5' to 3'): *RIPK1* Forward: CGTGGGAGTGATGTGTTGGA, *RIPK1* Reverse: ATCCTCCTGCCAAAAGTGCT, *CFLAR* Forward: ATGAACTTGTCTGGTTTGCAG, *CFLAR* Reverse: GCCTGCTTCCCTCTCTCTGTA. All clones were sequence-validated and both alleles of each clone contained mutations which resulted in premature stop codons.

##### Generation of *mCherry*- and *CFLAR*-overexpressing THP-1 cell lines

Human *CFLAR* was cloned into the pTwist Lenti SFFV Puro WPRE lentiviral vector backbone (Twist Bioscience). As a negative control, *mCherry* was cloned into pTwist Lenti SFFV Puro WPRE vector with a C-terminal Streptag II (Twist Bioscience) (80). Production of lentiviral particles, transduction, and selection of target cells were performed as previously described (38).

**qRT-PCR**

THP-1 cells were replated and differentiated as described above in a 48-well tissue culture-treated plate at a concentration of $2\times10^5$ cells/well. The next day, RNA was harvested (2 wells/condition) and isolated with RNeasy kit according to manufacturer instructions (Qiagen). cDNA synthesis was performed using High-Capacity cDNA Reverse Transcription Kit (Thermo Fisher). qPCR was performed using SYBR Green SuperMix (VWR International) on a QuantStudio Flex6000 (Thermo Fisher). The following primer sequences were used (all 5' to 3'):

*RIPK1* Forward: TTACATGGAAAAGGCGTGATACA

*RIPK1* Reverse: AGGTCTGCGATCTTAATGTGGA

*CFLAR* Forward: ATTTGCCTGTATGCCCGAGC

*CFLAR* Reverse: CCTGAGTGAGTCTGATCCACAC

**Statistical Analysis**

Data were graphed and analyzed using GraphPad Prism 9 (San Diego, CA, USA) and presented as mean values + SD. Mean values were compared across conditions and P values were determined using one- or two-way analysis of variance (ANOVA) or t test, as indicated.

**Supplementary Figure Legends**

**Fig. S1. Characterizing *Yersinia*-induced cell death in human macrophages.** Cells were pre-treated with Nec-1 and infected with the following strains of *Yersinia*: *WT Y. pseudotuberculosis* (*Yptb)*, *ΔyopJ*, *WT Y. enterocolitica (Ye)*, or *ΔyopP*. Cytotoxicity was measured by LDH release. **(A)** BMDMs were infected for 4-6 h at MOI 20. **(B-C)** hMDMs were infected for 16-22 h. Each data point represents the mean of triplicate wells for each of 7-12 different human donor hMDMs. **(D)** Immunoblot analysis was performed on hMDM lysates 5 h after infection for caspase-9 and β-actin. Representative of two independent experiments with different human donors. ns, not significant, *p < 0.05, **p < 0.01, ***p < 0.001, ****p < 0.0001 by Tukey's multiple comparisons test. Graphs depict mean + SD.

**Fig. S2. Characterizing cell death signaling downstream of IKKα/β blockade in human macrophages. (A-E)** Cells were pre-treated with IKKi, Nec-1, IETD, QVD, GSK'872, and/or ZVAD and stimulated with LPS. **(A-B)** hMDM cytotoxicity was measured by LDH release after 10 h of stimulation. Each data point represents the mean of triplicate wells for each of 3 different human donors. **(C-D)** THP-1 cell cytotoxicity was measured by LDH release after 17-24 h of stimulation. **(E)** Immunoblot analysis was performed on hMDM lysates for caspase-9 and β-Actin. Representative of 2 independent experiments. **(F)** Depending on condition, THP-1 cells were LPS-primed ($L_{pr}$), pre-treated with IKKi, Nec-1, and/or MCC950, and stimulated with LPS or Nigericin, or infected with WT *Ye* for 23-24 h. IL-18 levels were measured by ELISA in the supernatant. N=3. ns, not significant, *$p < 0.05$, **$p < 0.01$, ***$p < 0.001$, ****$p < 0.0001$ by Tukey's multiple comparisons test. Graphs depict mean + SD.

**Fig. S3. Characterization of *RIPK1*$^{-/-}$ THP-1 cells.** 2 independent *RIPK1*$^{-/-}$ single-cell clones were generated with CRISPR-Cas9. **(A)** Schematic representation of the *RIPK1* gene with exons (arrows). gRNA target sequence is highlighted in pink text. **(B)** Sequence alignments of WT THP-1 and *RIPK1*$^{-/-}$ Clones 1 and 2 are shown for both alleles. Red highlighting represents the mutated region. **(C)** Immunoblot analysis was performed on WT and *RIPK1*$^{-/-}$ THP-1 cell lysates for RIPK1 and β-actin. **(D)** RT-qPCR was performed on WT and *RIPK1*$^{-/-}$ THP-1 cell lysates for RIPK1 expression relative to HPRT. **(E)** WT and *RIPK1*$^{-/-}$ THP-1 cells were stimulated with LPS for 18-24 h. TNF levels were measured in the supernatant by ELISA. ns, not significant, ***$p < 0.001$ by Dunnett's multiple comparisons test. Graphs depict mean + SD.

**Fig. S4. Characterization of *CFLAR*$^{-/-}$ and plenti-*CFLAR* THP-1 cells. (A-C)** 2 independent *CFLAR*$^{-/-}$ single-cell clones were generated with CRISPR-Cas9. **(A)** Schematic representation of the *CFLAR* gene with exons (arrows). gRNA target sequence is highlighted in pink text. **(B)** Sequence alignments of WT THP-1 and *CFLAR*$^{-/-}$ Clones 1 and 2 are shown for both alleles. Red highlighting represents the mutated region. **(C)** Immunoblot analysis was performed on lysates for cFLIP and β-actin. **(D)** Immunoblot analysis was performed for cFLIP on lysates from plenti-*mCherry* and plenti-*CFLAR* stably-overexpressing THP-1 cells.

**Fig. S5. Jurkat and PANC-1 cells undergo RIPK1-dependent cell death following cIAP1/2 blockade.** Cells were pre-treated with Smac Mimetic ± Nec-1 and stimulated with TNF. Viability was measured by ATP signal in the following cell types: **(A)** THP-1 cells, 22-25 h, N=4 , **(B)** hMDMs, 5-6 h, N=3 , **(C)** Jurkat cells, 12-15 h, N=4, **(D)** PANC-1 cells, 14-16 h, N=4. ns, not significant, *p < 0.05, **p < 0.01, ***p < 0.001, ****p < 0.0001 by Tukey's multiple comparisons test. Graphs depict mean + SD.

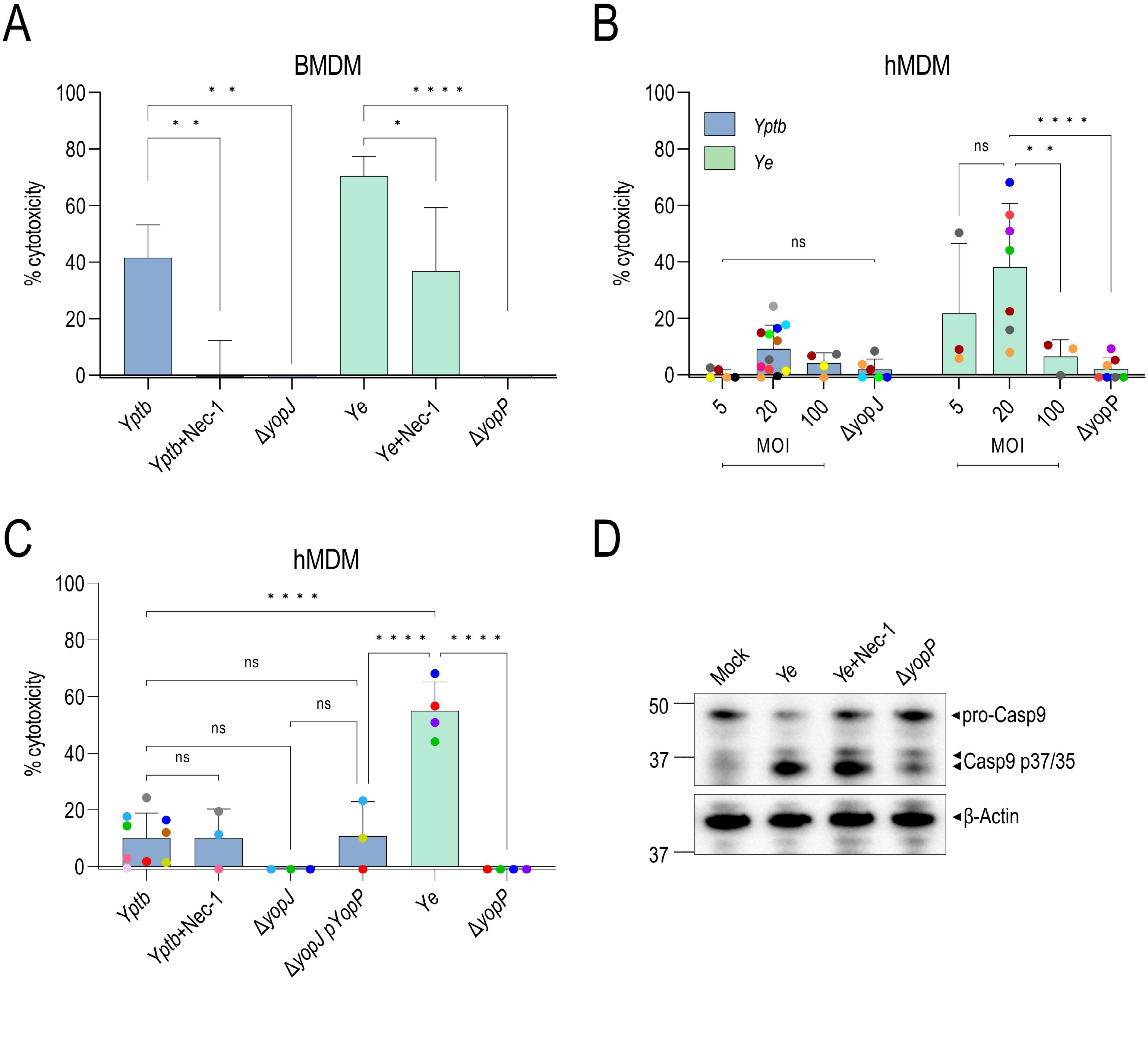
A
BMDM
% cytotoxicity
100
80
60
40
20
0
* *
* *
* * * *
*
Yptb
Yptb+Nec-1
ΔyopJ
Ye
Ye+Nec-1
ΔyopP
B
hMDM
% cytotoxicity
Yptb
Ye
ns
ns
* *
* * * *
5
20
100
ΔyopJ
5
20
100
ΔyopP
MOI
MOI
C
hMDM
% cytotoxicity
* * * *
* * * *
* * * *
ns
ns
ns
ns
Yptb
Yptb+Nec-1
ΔyopJ
ΔyopJ pYopP
Ye
ΔyopP
D
Mock
Ye
Ye+Nec-1
ΔyopP
50
37
37
pro-Casp9
Casp9 p37/35
β-Actin

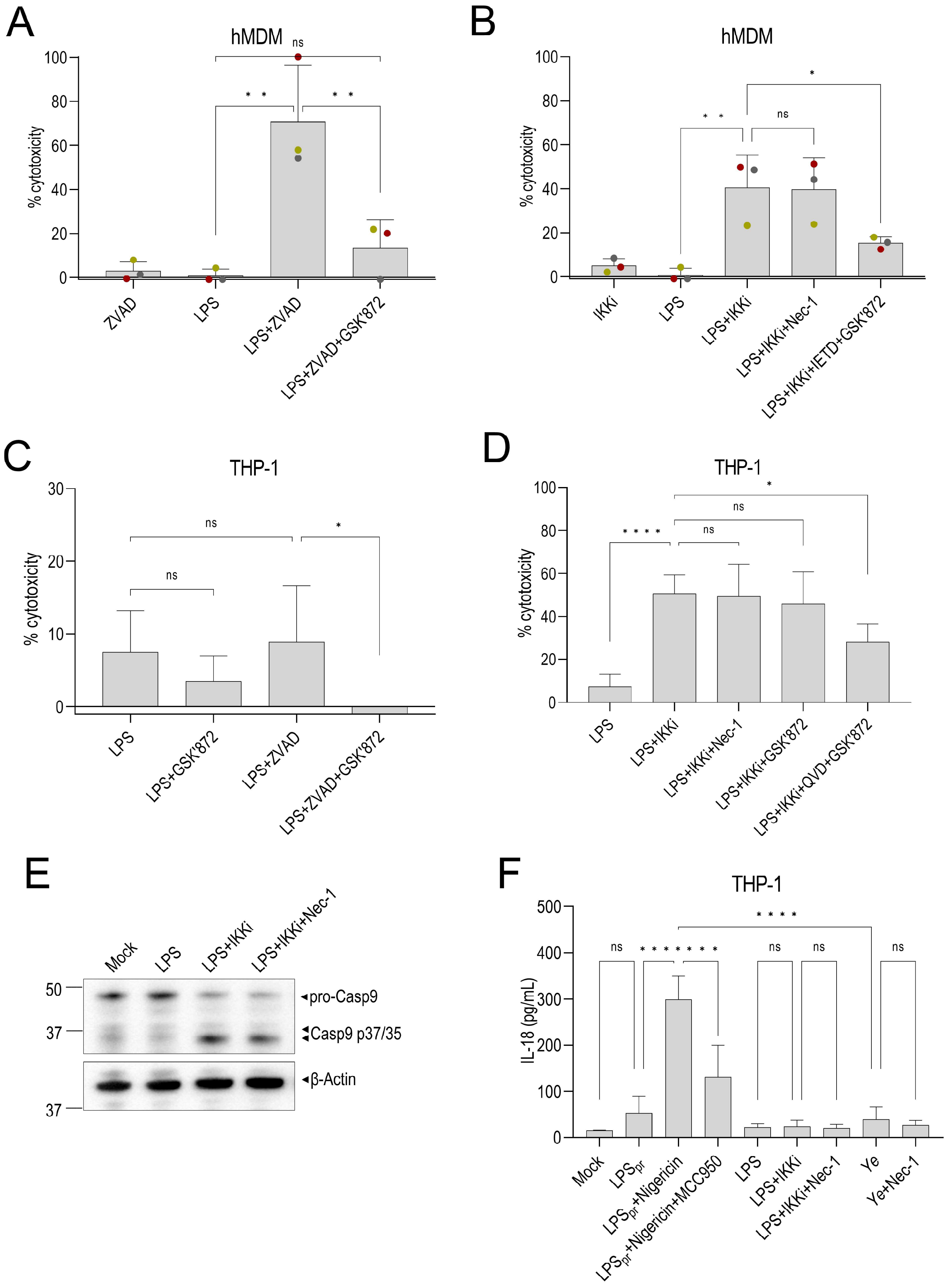
A
hMDM
% cytotoxicity
ZVAD
LPS
LPS+ZVAD
LPS+ZVAD+GSK'872
B
hMDM
% cytotoxicity
IKKi
LPS
LPS+IKKi
LPS+IKKi+Nec-1
LPS+IKKi+IETD+GSK'872
C
THP-1
% cytotoxicity
LPS
LPS+GSK'872
LPS+ZVAD
LPS+ZVAD+GSK'872
D
THP-1
% cytotoxicity
LPS
LPS+IKKi
LPS+IKKi+Nec-1
LPS+IKKi+GSK'872
LPS+IKKi+QVD+GSK'872
E
Mock
LPS
LPS+IKKi
LPS+IKKi+Nec-1
pro-Casp9
Casp9 p37/35
β-Actin
F
THP-1
IL-18 (pg/mL)
Mock
LPSpr
LPSpr+Nigericin
LPSpr+Nigericin+MCC950
LPS
LPS+IKKi
LPS+IKKi+Nec-1
Ye
Ye+Nec-1

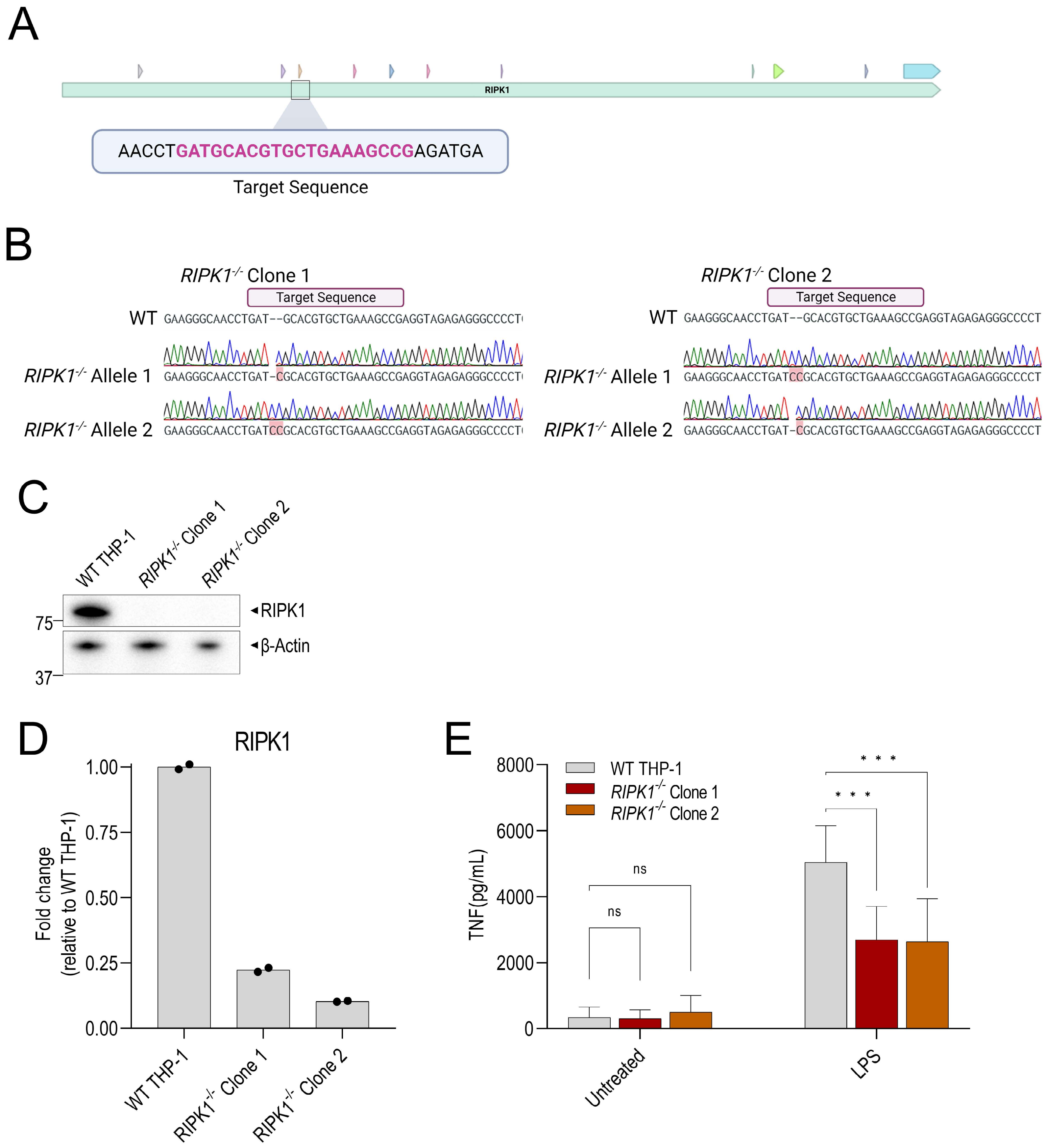
A
RIPK1
AACCTGATGCACGTGCTGAAAGCCGAGATGA
Target Sequence
B
RIPK1-/- Clone 1
RIPK1-/- Clone 2
Target Sequence
WT GAAGGGCAACCTGAT--GCACGTGCTGAAAGCCGAGGTAGAGAGGGCCCCT
RIPK1-/- Allele 1
RIPK1-/- Allele 2
C
WT THP-1
RIPK1-/- Clone 1
RIPK1-/- Clone 2
RIPK1
β-Actin
75
37
D
RIPK1
Fold change (relative to WT THP-1)
1.00
0.75
0.50
0.25
0.00
E
WT THP-1
RIPK1-/- Clone 1
RIPK1-/- Clone 2
TNF(pg/mL)
8000
6000
4000
2000
0
ns
* * *
Untreated
LPS

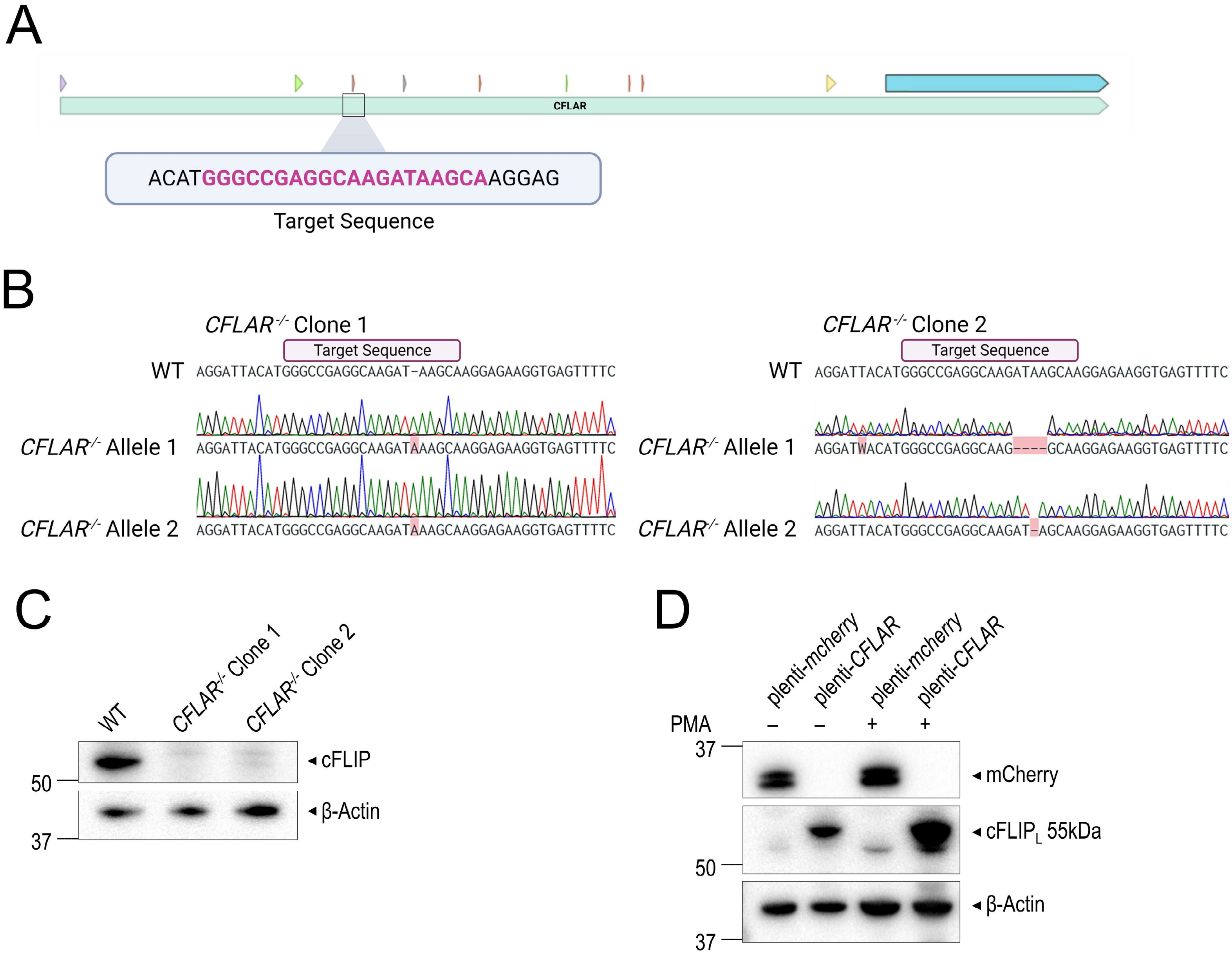
A
CFLAR
ACATGGGCCGAGGCAAGATAAGCAAGGAG
Target Sequence
B
CFLAR-/- Clone 1
Target Sequence
WT AGGATTACATGGGCCGAGGCAAGAT-AAGCAAGGAGAAGGTGAGTTTTC
CFLAR-/- Allele 1 AGGATTACATGGGCCGAGGCAAGATAAAGCAAGGAGAAGGTGAGTTTTC
CFLAR-/- Allele 2 AGGATTACATGGGCCGAGGCAAGATAAAGCAAGGAGAAGGTGAGTTTTC
CFLAR-/- Clone 2
Target Sequence
WT AGGATTACATGGGCCGAGGCAAGATAAGCAAGGAGAAGGTGAGTTTTC
CFLAR-/- Allele 1 AGGATWACATGGGCCGAGGCAAG----GCAAGGAGAAGGTGAGTTTTC
CFLAR-/- Allele 2 AGGATTACATGGGCCGAGGCAAGAT-AGCAAGGAGAAGGTGAGTTTTC
C
WT
CFLAR-/- Clone 1
CFLAR-/- Clone 2
cFLIP
β-Actin
50
37
D
plenti-mcherry
plenti-CFLAR
plenti-mcherry
plenti-CFLAR
PMA – – + +
37
mCherry
cFLIPL 55kDa
50
β-Actin
37

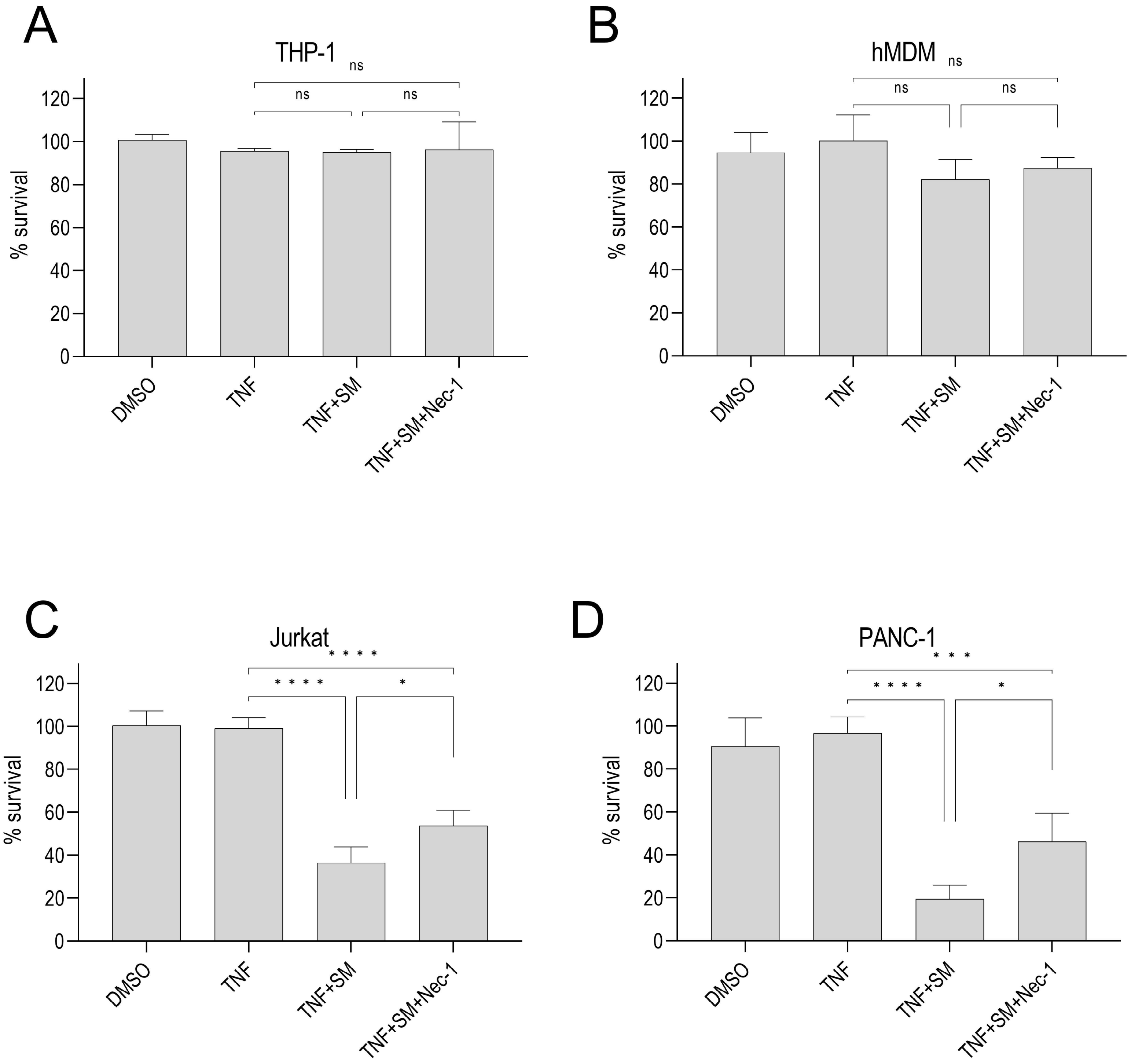
A
THP-1
ns
ns
ns
% survival
0
20
40
60
80
100
120
DMSO
TNF
TNF+SM
TNF+SM+Nec-1
B
hMDM
ns
ns
ns
% survival
0
20
40
60
80
100
120
DMSO
TNF
TNF+SM
TNF+SM+Nec-1
C
Jurkat
* * * *
* * * *
*
% survival
0
20
40
60
80
100
120
DMSO
TNF
TNF+SM
TNF+SM+Nec-1
D
PANC-1
* * *
* * * *
*
% survival
0
20
40
60
80
100
120
DMSO
TNF
TNF+SM
TNF+SM+Nec-1

### Relating to Figures 1D, S1D, 2C, S2E

Antibody: Casp8

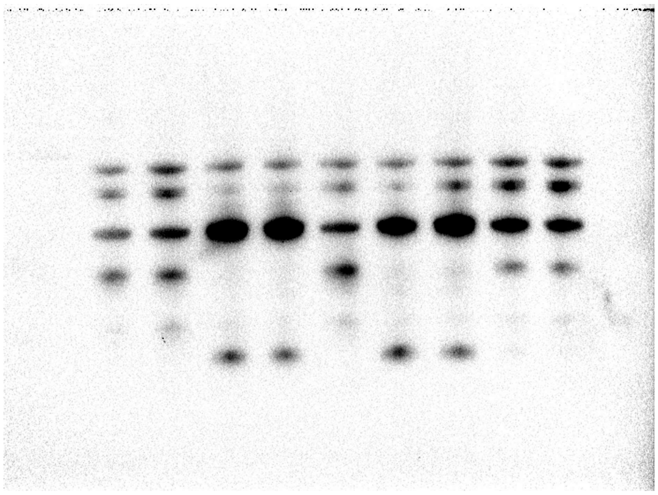

Antibody: cleaved Casp8

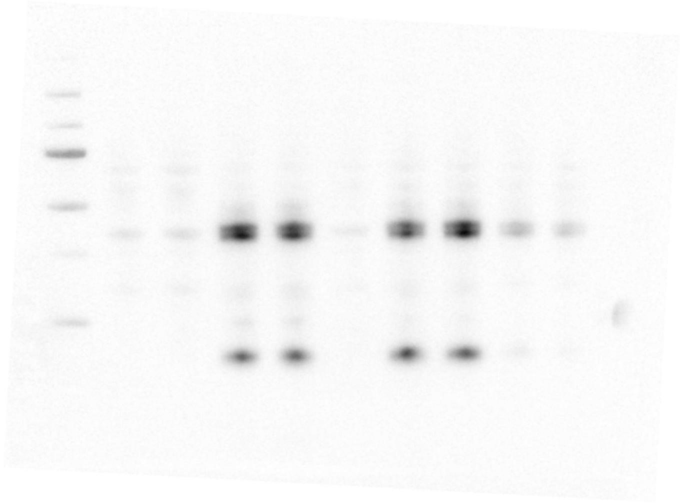

Antibody: Casp9

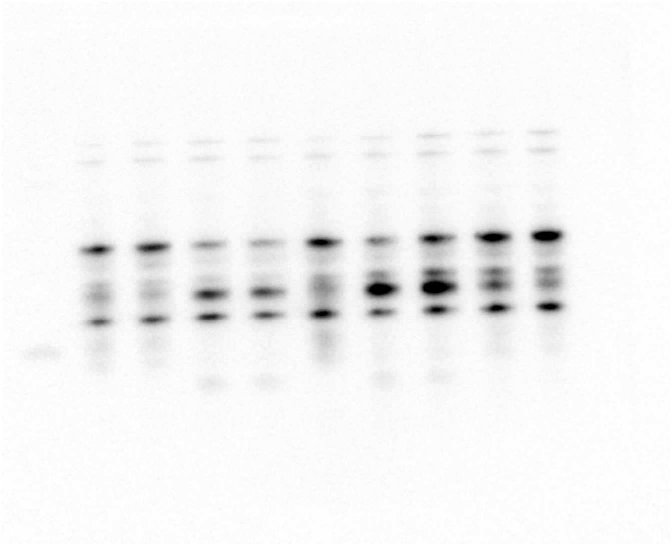

Antibody: β-Actin

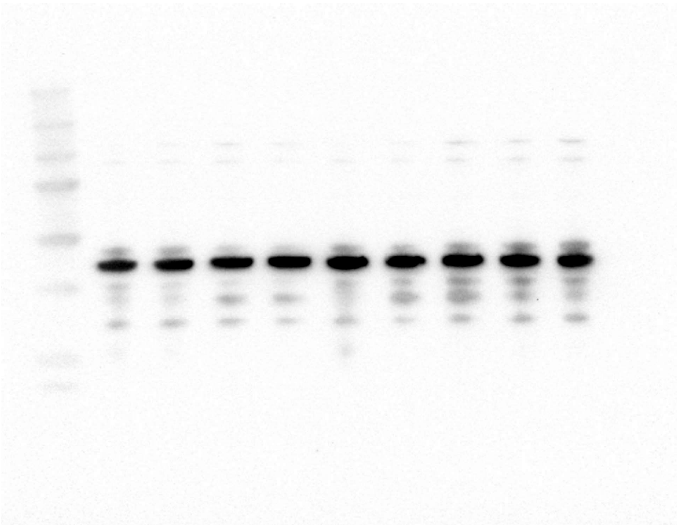

Ladder (Colorimetric)

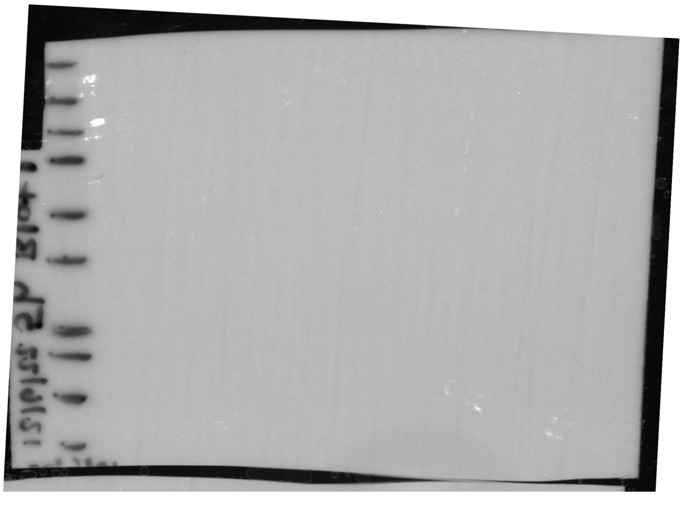

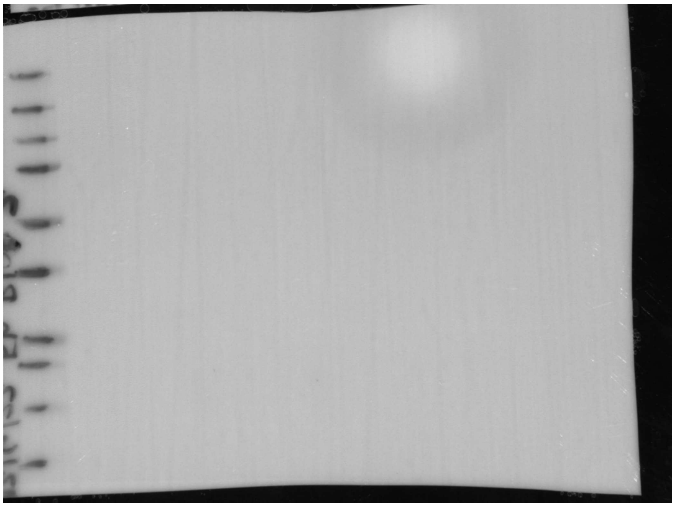

### Relating to Figure 1E

Antibody: Casp3

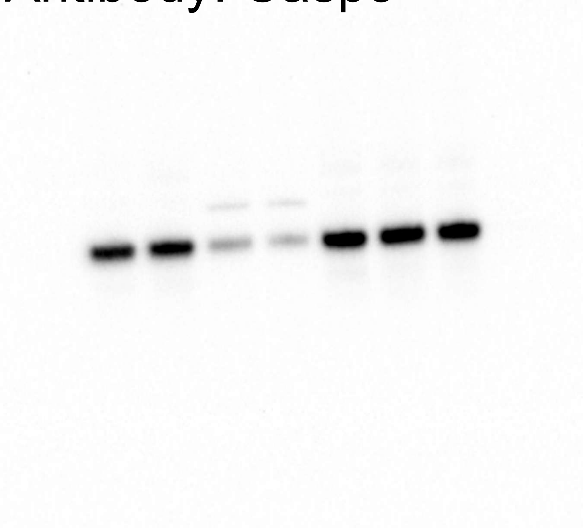

Antibody: cleaved Casp3

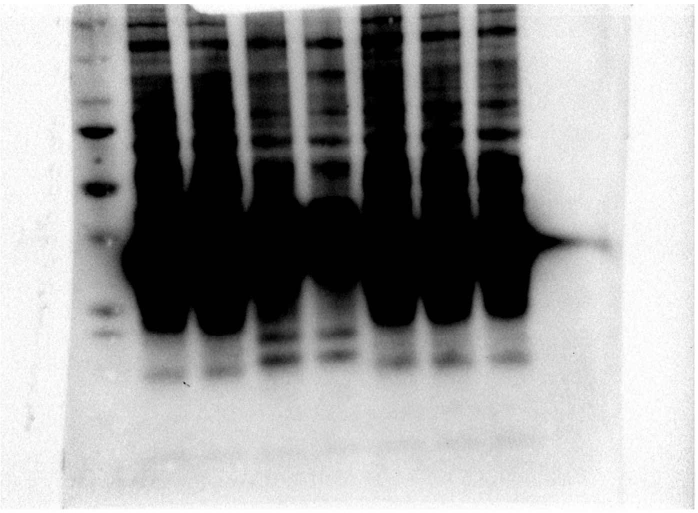

Antibody: β-Actin

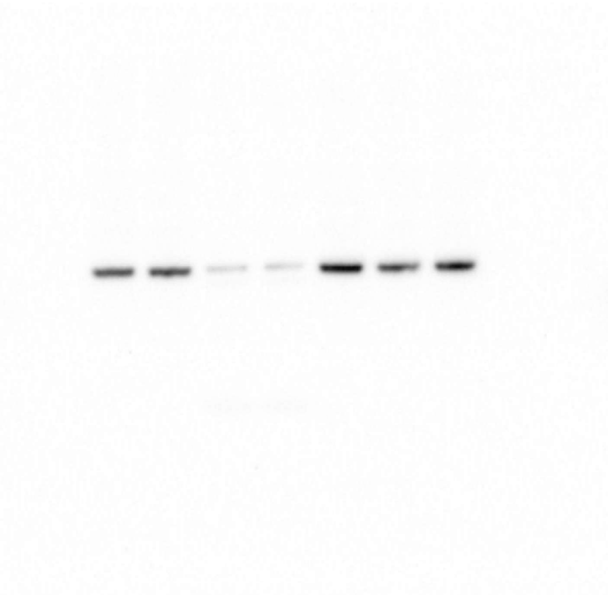

Ladder (Colorimetric)

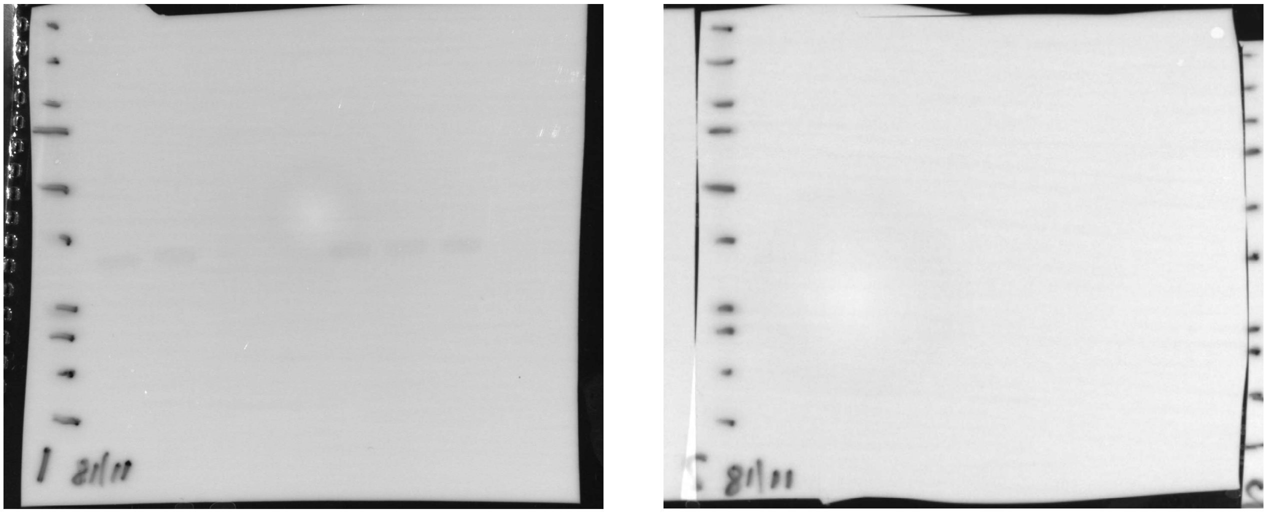

### Relating to Figure 1G

#### Antibody: Casp3

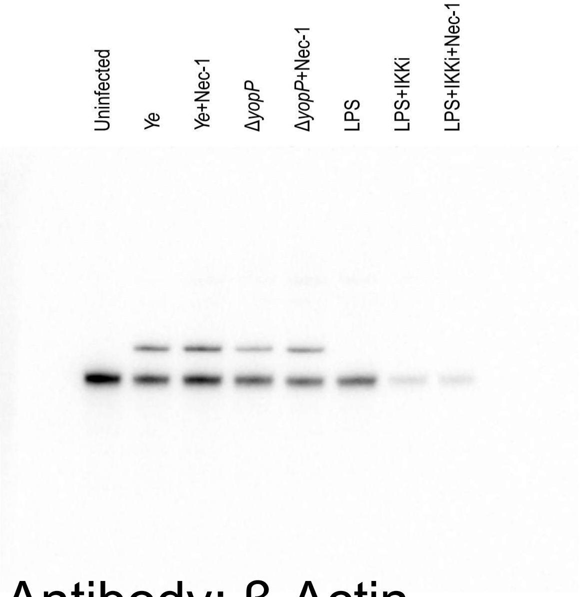


#### Antibody: cleaved Casp3

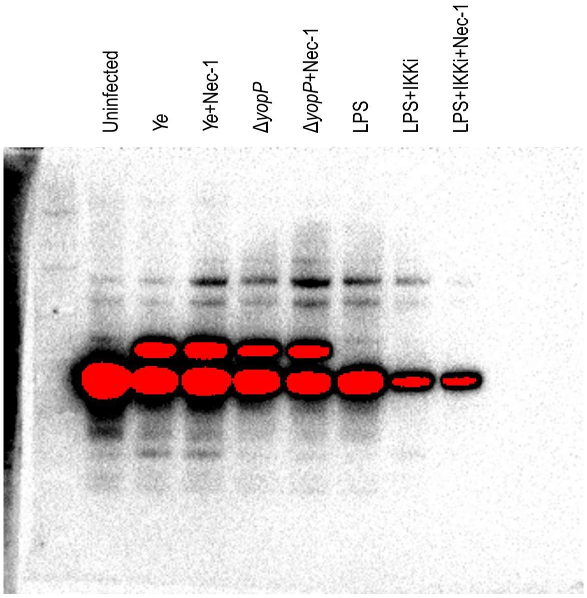


#### Antibody: β-Actin

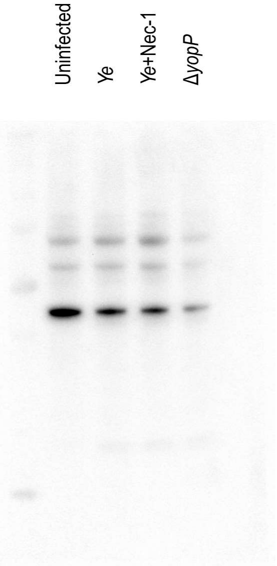


#### Ladder (Colorimetric)

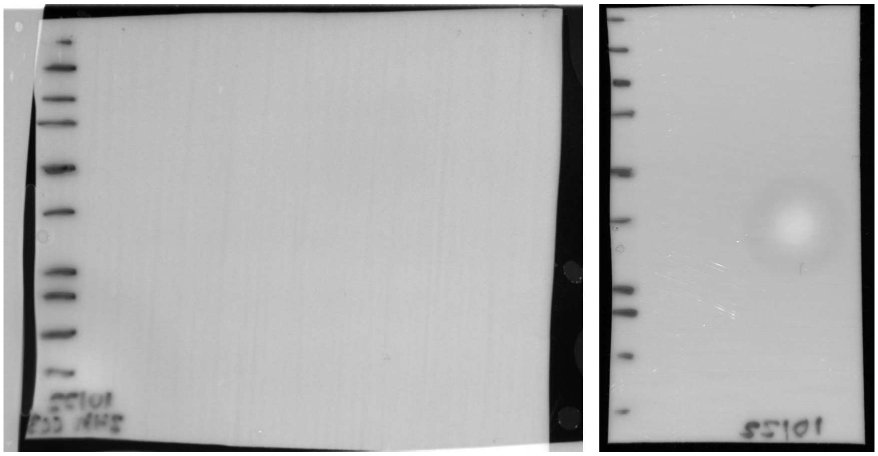

### Relating to Figure 2D

Antibody: RIPK1

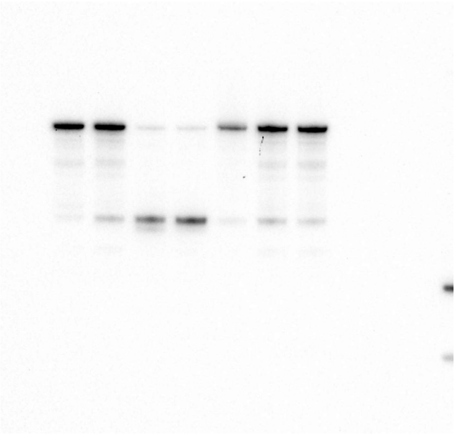

Antibody: p-RIPK1 (S166)

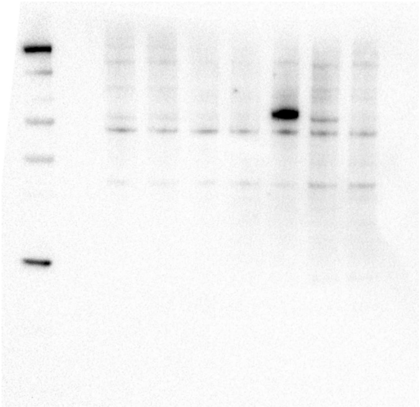

Antibody: Casp3

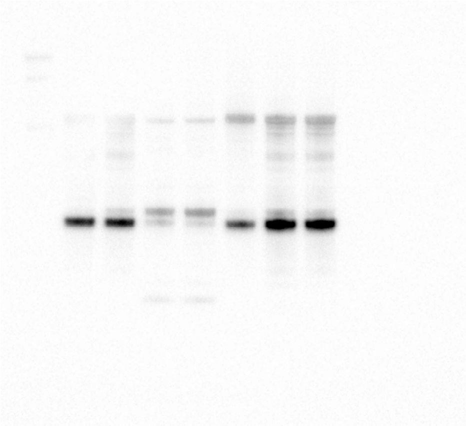

Antibody: cleaved Casp3

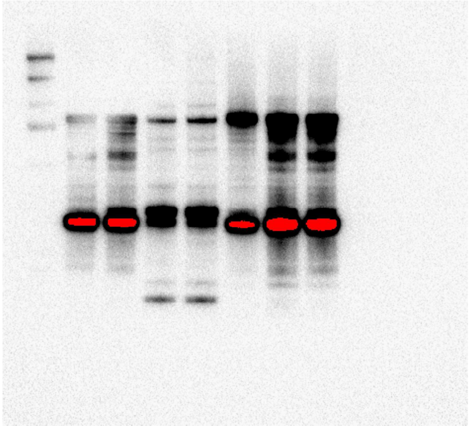

Antibody: β-Actin

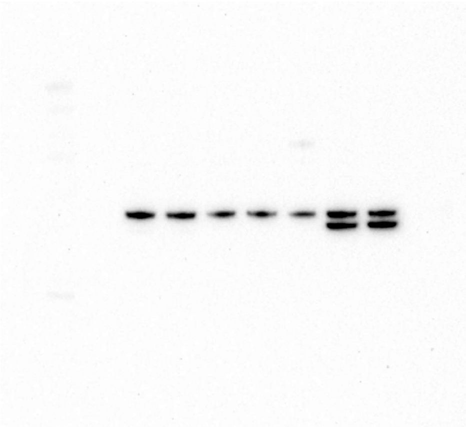

Ladder (Colorimetric)

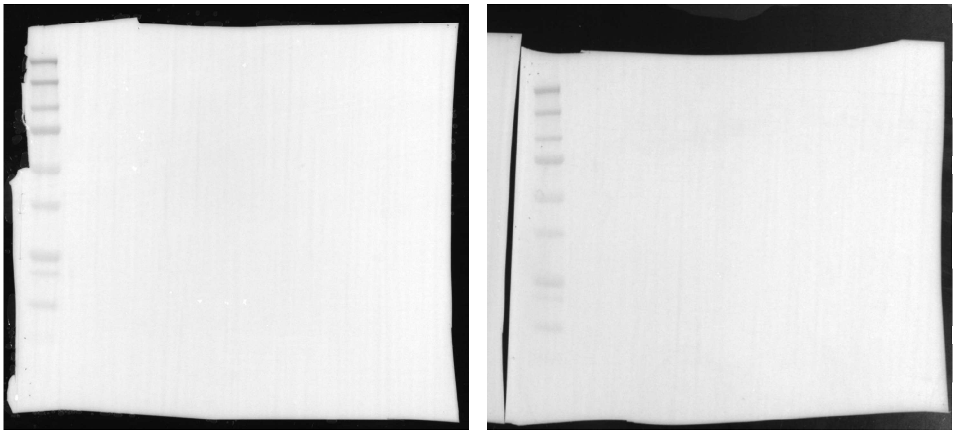

### Relating to Figure 2F

Antibody: Casp3

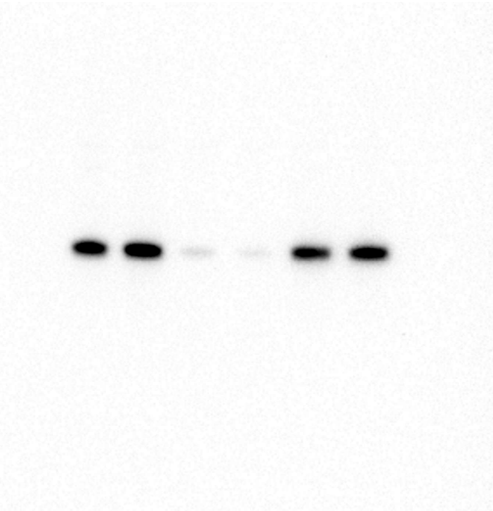

Antibody: cleaved Casp3

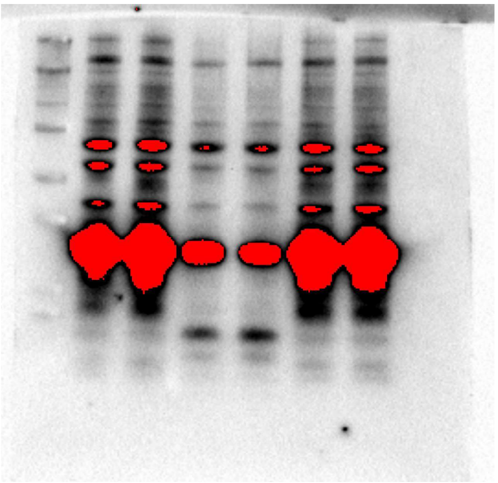

Antibody: β-Actin

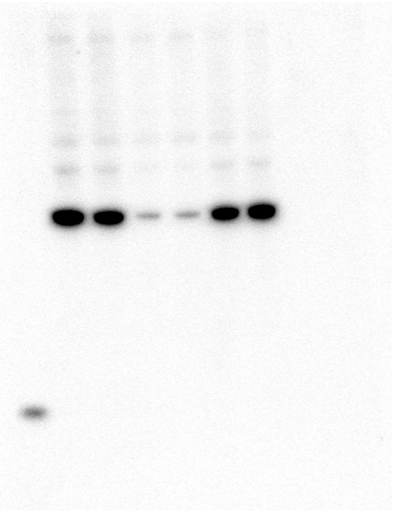

Ladder (Colorimetric)

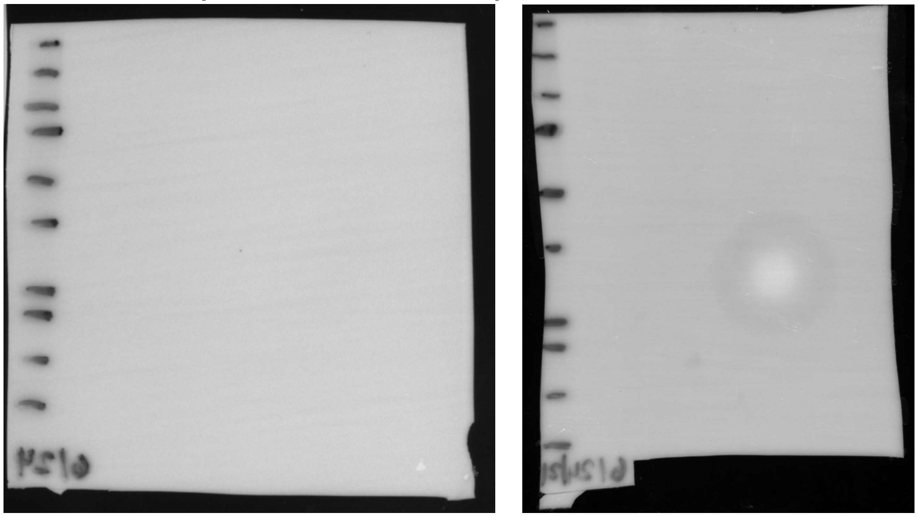

### Relating to Figure S3C

Antibody: RIPK1

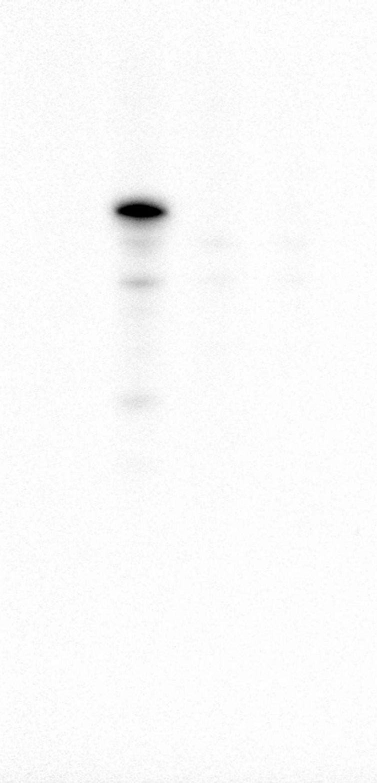

Antibody: β-Actin

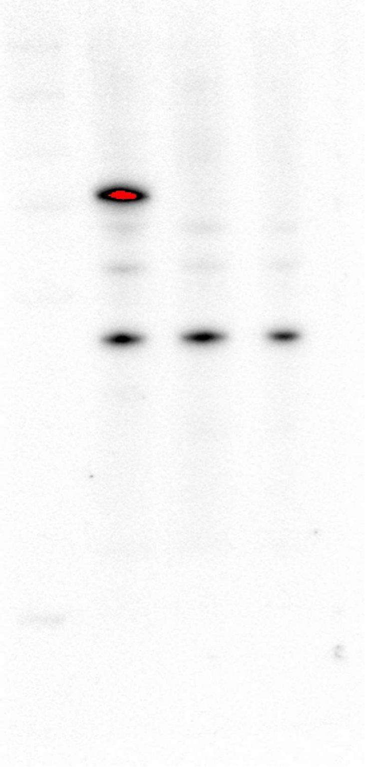

Ladder (Colorimetric)

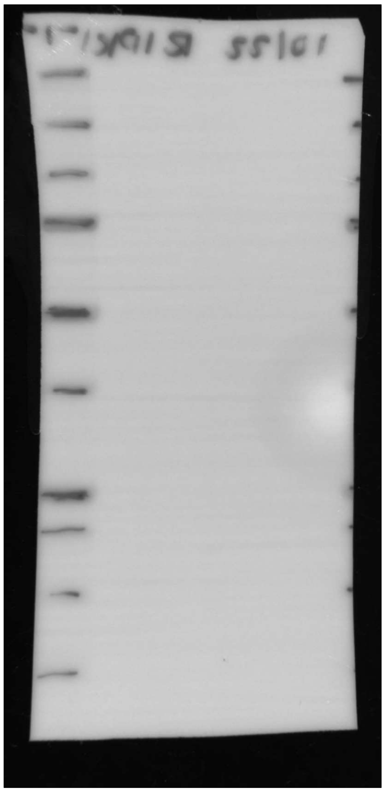

### Relating to Figure 4A

Antibody: CFLIP

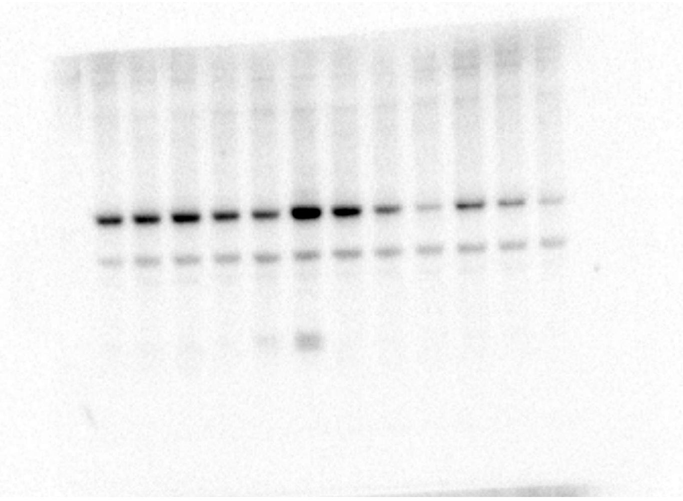

Antibody: β-Actin

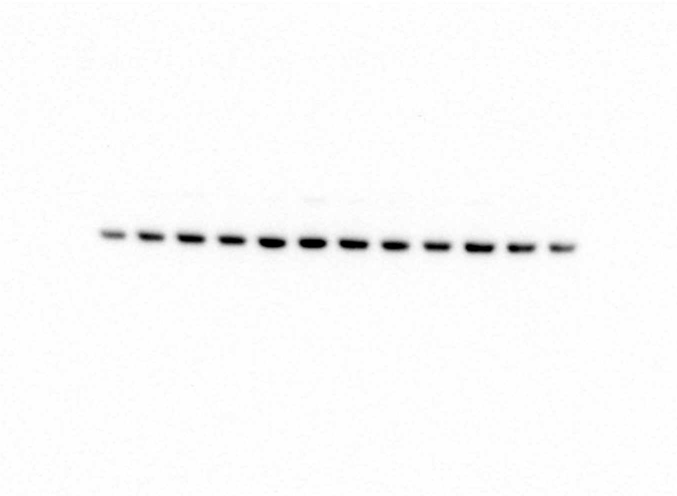

Ladder (Colorimetric)

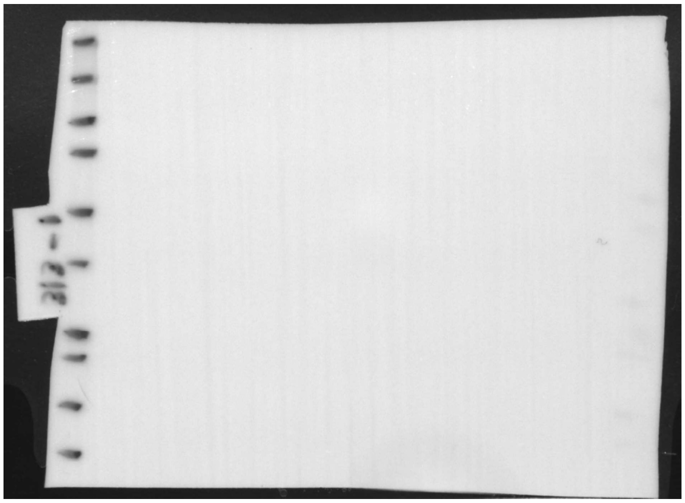

### Relating to Figure S4C

Antibody: CFLIP

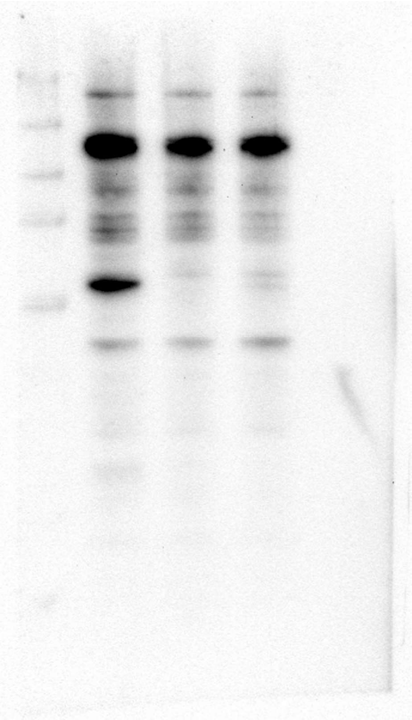

Antibody: β-Actin

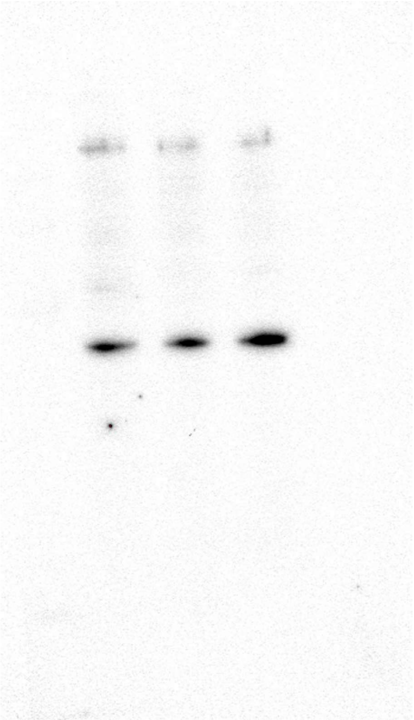

Ladder (Colorimetric)

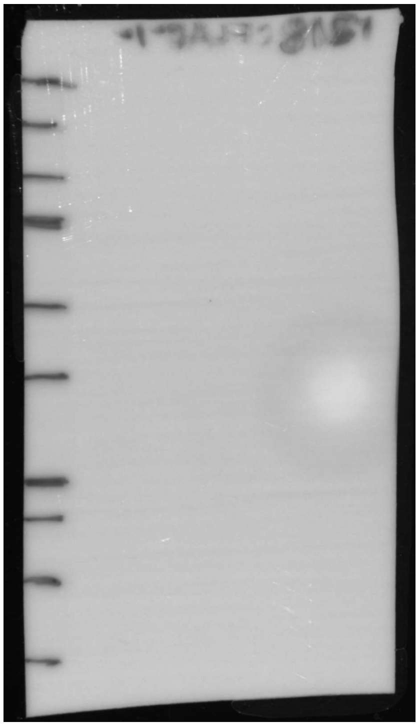

### Relating to Figure S4D

Antibody: CFLIP

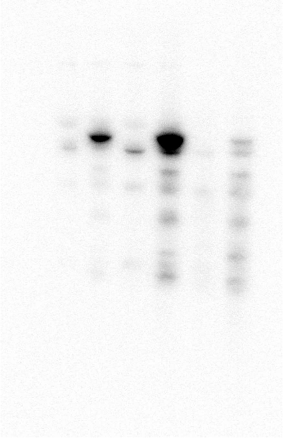

Antibody: mCherry

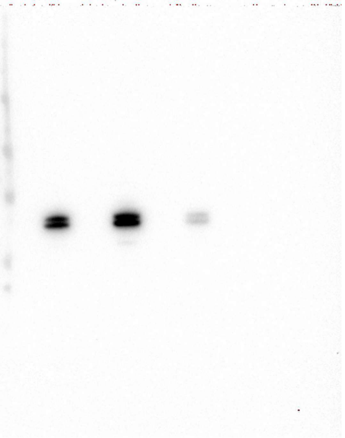

Antibody: β-Actin

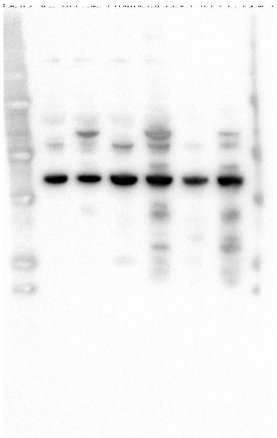

Ladder (Colorimetric)

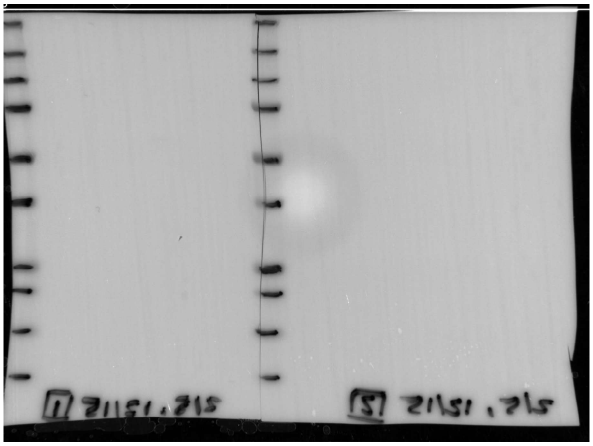
